# Diversity and prevalence of colibactin- and yersiniabactin encoding mobile genetic elements in enterobacterial populations: insights into evolution and co-existence of two bacterial secondary metabolite determinants

**DOI:** 10.1101/2021.01.22.427840

**Authors:** Haleluya Wami, Alexander Wallenstein, Daniel Sauer, Monika Stoll, Rudolf von Bünau, Eric Oswald, Rolf Müller, Ulrich Dobrindt

## Abstract

The bacterial genotoxin colibactin interferes with the eukaryotic cell cycle by causing double-stranded DNA breaks. It has been linked to bacterially induced colorectal cancer in humans. Colibactin is encoded by a 54-kb genomic region in *Enterobacteriaceae*. The colibactin genes commonly co-occur with the yersiniabactin biosynthetic determinant. Investigating the prevalence and sequence diversity of the colibactin determinant and its linkage to the yersiniabactin operon in prokaryotic genomes, we discovered mainly species-specific lineages of the colibactin determinant and classified three main structural settings of the colibactin-yersiniabactin genomic region in *Enterobacteriaceae*. The colibactin gene cluster has a similar but not identical evolutionary track to that of the yersiniabactin operon. Both determinants could have been acquired on several occasions and/or exchanged independently between enterobacteria by horizontal gene transfer. Integrative and conjugative elements play(ed) a central role in the evolution and structural diversity of the colibactin-yersiniabactin genomic region. Addition of an activating and regulating module (*clbAR*) to the biosynthesis and transport module (*clbB-S*) represents the most recent step in the evolution of the colibactin determinant. In a first attempt to correlate colibactin expression with individual lineages of colibactin determinants and different bacterial genetic backgrounds, we compared colibactin expression of selected enterobacterial isolates *in vitro*. Colibactin production in the tested *Klebsiella* spp. and *Citrobacter koseri* strains was more homogeneous and generally higher than that in most of the *E. coli* isolates studied. Our results improve the understanding of the diversity of colibactin determinants and its expression level, and may contribute to risk assessment of colibactin-producing enterobacteria.

## 2 Introduction

The non-ribosomal peptide/polyketide hybrid colibactin is a secondary metabolite found in a variety of bacterial species of the family *Enterobacteriaceae*. The colibactin biosynthetic machinery is encoded by a 54-kb large polyketide synthase (*pks*) or *clb* genomic island[1], which includes 19 genes. The largest part of the island consists of a section of overlapping or closely spaced genes: *clbB* to *clbL* and *clbN* to *clbQ*, which are aligned on the same strand and code for components of the biosynthesis complex. The colibactin assembly line is supplemented with a dedicated transporter, encoded by *clbM*, and a resistance-conferring protein encoded by *clbS* [2, 3]. Two additional genes required for colibactin production are located ca. 400 bp upstream of the first biosynthesis gene *clbB* in the opposing reading direction: the *clbR* gene coding for an auto-activating, *pks* island-specific transcription factor and the phosphopantetheinyl transferase-encoding gene *clbA*, which is crucial for activation of polyketide biosynthesis complexes (Fig. 1) [4–6]. Between these two divergent transcription units, there is a “variable number of tandem repeat” (VNTR) region, which comprises a varying number of a repeating octanucleotide sequence (5’-ACAGATAC-3’) depending on the isolate [2].

**Figure 1.**
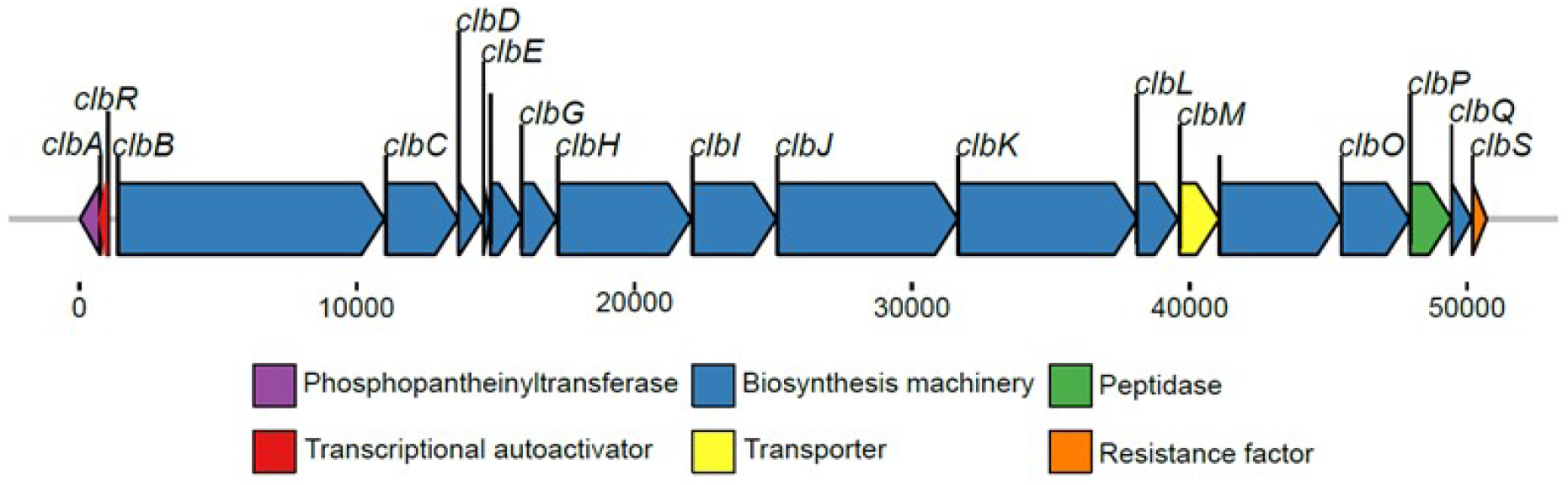
Schematic representation of the genomic architecture of the *pks* island (ca. 54 kb) present in *E. coli* strain M1/5. The 19 genes within the island are colored with respect to their function. The island codes for a phosphopantetheinyl transferase (*clbA*, in purple), a transcriptional autoactivator (*clbR*, in red), multiple core biosynthetic genes (*clbB-clbL, clbN, clbO*, and *clbQ* in blue), a transporter (*clbM*, yellow), a peptidase (*clbP*, in green) and a resistance factor (*clbS*, orange).

Recently, the structure of the colibactin molecule has been proposed [3, 7–9]. Yet, the biological role of colibactin is still under discussion. Colibactin can interfere with the progression of the eukaryotic cell cycle, presumably by cross-linking DNA resulting in double-strand DNA breaks in genomic instability in eukaryotes [1, 10, 11]. The ability to produce colibactin has been described to increase the pathogenic potential of the producing bacteria and to promote colorectal cancer development [12–17], but has also been related to beneficial effects to the host [3, 18–20].

Initially, the *pks* island has been described in extraintestinal pathogenic *E. coli* to be chromosomally inserted into the *asnW* tRNA locus in close proximity to another tRNA(*Asn*) gene-associated pathogenicity island, the so-called “high pathogenicity island” (HPI). The HPI harbors an additional polyketide determinant coding for the metallophore yersiniabactin biosynthetic machinery [1, 21]. As members of the *Enterobacteriaceae* are generally not known as archetypal secondary metabolite producers the origin of the *pks* island remains to be further investigated. Interestingly, the colibactin determinant has also been detected as part of an “integrative and conjugative element” (ICE) in different enterobacteria. This ICE also integrates near a tRNA(*Asn*) locus into the bacterial chromosome and commonly carries the yersiniabactin gene cluster [2, 22]. It has been suggested that the close linkage observed between the colibactin and yersiniabactin gene clusters results from the functional interconnection between the colibactin and yersiniabactin biosynthetic pathways via the phosphopantetheinyl transferase ClbA, which can also contribute to the biosynthesis of yersiniabactin [5, 23]. The highly conserved colibactin determinant has so far been detected in a spectrum of strains belonging to the *Enterobacteriaceae* family: most commonly among *E. coli* strains of the phylogroup B2, followed by *Klebsiella pneumoniae* isolates, but also in *Citrobacter koseri* and *Klebsiella aerogenes* [2, 18, 24]. Less conserved variants or homologs of the colibactin gene cluster have been described phenotypically or based on nucleotide sequence data in an *Erwinia oleae* strain, the honey bee symbiont *Frischella perrara*, and the marine α-proteobacterium *Pseudovibrio* [24, 25].

Based on the low sequence similarity between the enterobacterial colibactin genes and the two homologous polyketide determinants in *F. perrara* and *Pseudovibrio* as well as on their association with mobile genetic elements (MGEs) or at least mobility-associated genes, one can hypothesize that the colibactin gene cluster is spread by horizontal gene transfer, maybe via an ICE-like element [2, 24]. While in most *Enterobacteriaceae* the colibactin determinant is typically associated with an ICE, the characteristic mobility and transfer features of an ICE are absent in the sequence context of the *pks* island in *E. coli* phylogroup B2 strains. Nevertheless, the pks island in E. coli remains mobilizable and transferable through external factors, supporting the hypothesis that former MGEs can undergo a stabilization (homing) process upon their chromosomal integration [26–28].

Studies addressing the prevalence of the colibactin genes were so far mainly focused on *Klebsiella* spp. or *Escherichia coli* backgrounds. The scarcity of data in other prokaryotic species regarding its distribution and the structure of the associated MGE makes it challenging to reliably further characterize the transmission and evolution of this polyketide determinant [18]. Previous data show that the prevalence varied from 5.3% to 25.6% in *Klebsiella* and from 9.5% to 58% in *Escherichia*, highlighting an enrichment of *pks* island in specific ecological niches, whereas studies with a broader screening approach resulted in a prevalence of 14% in *Klebsiella* and 9.5% in *Escherichia* isolates, respectively [2, 29–34]. Notably, in health-related studies, a higher association of the *clb* genes was observed amongst strains with an increased virulence potential, with a prevalence of as much as 78.8% for *Klebsiella* subgroups and 72.7% for colorectal cancer-associated *E. coli* isolates [29, 32, 35–37]. The colibactin genes are frequently found in hyper-virulent and multidrug-resistant *K. pneumoniae* isolates [38, 39]. The obvious prevalence of the colibactin gene cluster in specific enterobacterial species combined with the description of more distantly related homologous determinants has sparked our interest in a better understanding of the spread and evolution of the colibactin determinant and its genetic context in bacteria. Therefore, our study aimed to investigate the prevalence and diversity of the colibactin determinant also in isolates outside of the *Enterobacteriaceae* family. Furthermore, we compared colibactin expression levels among enterobacterial isolates carrying different lineages of colibactin determinants as a first attempt to assess the functional context of the bacterial genetic background, pathogenicity, and colibactin expression.

## 3 Impact Statement

Colibactin can act as a bacterial genotoxin and thus promote colorectal cancer development. Little is known about the origin, diversity and prevalence of the colibactin genes (*clb*) within prokaryotes. The *clb* genes are closely associated with pathogenicity islands or integrative and conjugative elements (ICEs). We screened roughly 375,000 prokaryotic genomes to analyze the diversity and evolution of such mobile genetic elements among bacterial populations. Interestingly, *clb* genes are only present in a subgroup of enterobacteria, mainly *E. coli*, *Klebsiella* spp. and *Citrobacter koseri*. The *clb* determinant, together with the yersiniabactin (*ybt*) gene cluster, belong to an ICE in most of the *clb*-positive enterobacteria, especially in *Klebsiella*. We show that both determinants, though in principle freely transferable within bacteria, have a mainly species-specific phylogeny, and that colibactin expression levels were species-independent. Recombination promoted the structural diversification of the ICE in different species, including its successive degeneration that led to the establishment of the colibactin and yersiniabactin islands in *E. coli* phylogroup B2 strains. Our results not only illustrate differing evolutionary tracks of the *clb* and *ybt* determinants in different enterobacterial species, but also highlight the important of ICEs for genomic variability in enterobacteria and the evolution of archetypal pathogenicity islands.

## 4 Methods

### 4.1 Bacterial strains and media

For cultivation, bacteria were grown as batch cultures in lysogeny broth (LB) (10g/l tryptone, 5g/l yeast extract, 5g/l sodium chloride) at 37°C. Strains used in this study are listed in the Supplemental Material (Table S1).

### 4.2 DNA extraction and sequencing

DNA extraction of the enterobacterial strains was performed using the MagAttract^®^ HMW DNA Kit (Qiagen, Hilden, Germany) according to the manufacturer’s recommendations. To prepare paired-end libraries we used the Nextera XT DNA Library Preparation kit (Illumina, San Diego, CA, USA). Libraries were sequenced on the Illumina MiSeq sequencing platform using v2 sequencing chemistry (500 cycles) or on the Illumina NextSeq500 system using v2.5 chemistry (300 cycles). Accession numbers of in-house sequences submitted to the NCBI GenBank database are included in Supplemental Material (Table S1).

### 4.3 Genome selection and phylogenetic analysis

All genome sequences not generated in this study were obtained from publically available prokaryotic genomes (NCBI GenBank). The quality of in-house sequenced genomes was checked with FastQC v0.11.5 (https://github.com/chgibb/FastQC0.11.5/blob/master/fastqc), and low-quality reads were trimmed using Sickle v1.33 (https://github.com/najoshi/sickle). The processed reads were *de novo* assembled with SPAdes v3.13.1 [40] and annotated with prokka v1.12 [41]. The genomes were screened for the presence of >45 kb of the complete *pks* genomic island using standalone BLAST+ v2.8.1 [42] and antiSMASH v5.0.0 [43]. The *pks* island found in the genome of the *E. coli* strain M1/5 (accession no. CP053296) was used as a reference sequence. The VNTR and the sequence stretch of the *pks* island that spans between *clbJ* and *clbK* were excluded from analysis as these regions are prone to misassembly.

The contigs that align to the colibactin genes were ordered using ABACAS v1.3.1 [44] and multiple sequence alignment was generated using Kalign v3.1.1 [45]. Recombinant regions were detected and removed using Gubbins v2.4.1 [46]. The recombination filtered polymorphisms were then used to generate a maximum likelihood phylogeny of the colibactin determinant using RAxML v8.2.11 [47] under the GTR-GAMMAX model from 9974 polymorphic sites. The branch support of the maximum likelihood tree was estimated by bootstrap analysis of 200 replicate trees. The homologous gene cluster found in *F. perrara* was used as an outgroup. The phylogeny of the corresponding *ybt* islands was generated with a similar approach. The generated trees were visualized using itoL (https://itol.embl.de).

### 4.4 Phylo-grouping of *pks*-positive strains

The *E. coli* and *Klebsiella* strains that harbored the colibactin gene cluster were allocated to their corresponding sequence types using mlst v2.16.1 (https://github.com/tseemann/mlst), which detects sequence types using the PubMLST typing schemes. The *Escherichia* strains were further classified into their phylogenetic lineages using the standalone tool, EzClermont v0.4.5 (https://github.com/nickp60/EzClermont).

The analysis of the diversity of the colibactin and yersiniabactin gene clusters involved virulence gene multi-locus sequence typing (MLST) for both polyketide determinants as previously described [38]. Briefly, the allele sequences of sixteen genes of the colibactin gene cluster (*clbACDEFGHILMNOPQR*) as well as of eleven genes of the yersiniabactin determinant (*fyuA, ybtE, ybtT, ybtU, irp1, irp2, ybtA, ybtP, ybtQ, ybtX, ybtS*) were extracted from the individual genomes and analyzed for allelic variations. Each observed combination of alleles was assigned a unique colibactin sequence type (CbST, listed in Table S5) or yersiniabactin sequence type (YbST, listed in Table S6).

### 4.5 Variable number tandem repeat (VNTR) detection

The VNTR copy number present within the colibactin determinant (upstream of *clbR*) was detected using the standalone version of tandem repeats finder v4.09 [48]. The VNTR copy number distribution was visualized using R v3.4.3 (https://www.r-project.org/index.html).

### 4.6 Detection of *E. coli* virulence markers for pathotyping

For pathotyping, the *clb*-positive *E. coli* strains were *in silico* screened for the presence of different *E. coli* pathotype marker genes using BLAST+ v2.8.1 (Supplemental Material, Tab. S). These genes were used as markers for *E. coli* pathotypes: enteroaggregative *E. coli* (EAEC), enterohemorrhagic *E. coli* (EHEC), enteropathogenic *E. coli* (EPEC), enterotoxigenic *E. coli* (ETEC), diffusely adhering *E. coli* (DAEC), uropathogenic *E. coli* (UPEC), and newborn meningitis-causing *E. coli* (NMEC).

### 4.7 Quantification of colibactin expression through N-myristoyl-D-asparagine

Following an approach described by Bian and colleagues [49], a collection of colibactin-producing strains of the main species harboring the colibactin determinant was characterized for their ability to produce colibactin under *in vitro* growth conditions. For this purpose, we quantified N-myristoyl-D-asparagine (N-Myr-D-asparagine) a byproduct during colibactin maturation. The amount of this intermediate extrapolates the resulting colibactin amount produced. After growing the bacteria for 24 h at 37°C in glass tubes in 5 ml LB supplemented with 200 μl of a water/XAD-16-resin slurry, bacterial cells were harvested by centrifugation. The pelleted bacteria-slurry mixes were sedimented, filtered, and dissolved three times in acetone with increasing volume (12ml, 100 ml, and finally 200ml). Afterward, the solvent was exchanged by rotary evaporation and replaced by 1.6 ml methanol. The sample was further concentrated by centrifugation (10 min, 15000 rpm at 4°C), followed by drying 1,5 ml of the solution in a vacuum centrifuge and subsequent resuspension in 50 μl methanol. 30 μl of these processed samples were measured by Ultra Performance Liquid Chromatography coupled to High-Resolution Mass Spectrometry (UPLC-HRMS) conducted on a Thermo Scientific Ultimate 3000 RS with a Waters Acquity BEH 100*2.1mm 1.7μm 130A column (eluent A: 0.1% formic acid in ddH2O, eluent B: 0.1% formic acid in acetonitrile), where a flow rate of 0.6ml/min followed by a Bruker Maxis II-4G, 150-2500 m/z and a scan rate of 2Hz was applied. To enable quantification of N-Myr-D-asparagine, we used 250 mM cinnarizine as internal standard and normalized peak areas based on the internal standard and the optical density (OD_600_) of the bacterial culture.

## 5 Results

### 5.1 Prevalence of colibactin determinant

Of the 374,754 publicly accessible prokaryotic genomes (as of 30.06.2019) that were screened for the presence of the colibactin gene cluster, 1,969 genomes carried this polyketide-encoding operon. An additional 200 *clb*-positive enterobacterial genomes determined in-house were added to the analysis (Tab. 1, Supplemental Material, Tab. S1). The *clb* gene cluster was detected in several enterobacterial species, most frequently in *E. coli* and *K. pneumoniae* isolates, but also to a lesser extent in *K. aerogenes*, *C. koseri*, *E. cloacae*, *E. hormaechei*, *K. michiganensis*, *S. marcescens*, and *E. oleae*. The colibactin determinant was, however, not detectable in 112,546 *Salmonella enterica* and 41 *S. bongori* genomes (Supplemental Material, Tab. S4), but in one out of eight genomes of unspecified *Salmonella* isolates. We did not detect the *clb* genes in 2,634 *Shigella* spp., 861 *Yersinia* spp., 677 *Serratia* spp., 186 *Proteus* spp. and 69 *Morganella* spp. genomes (Supplemental Material, Tab. S4). A less well-conserved homolog of the colibactin determinant was detected in three *F. perrara* genomes. It should be noted that the number of genomes of *Klebsiella* spp. and *E. coli* analyzed in this study are markedly higher than those of the other species and lineages due to the sequencing bias towards *Klebsiella* spp. and *E. coli* strains. Accordingly, a reliable statement on the prevalence of the colibactin determinant in the different species cannot be made.

### 5.2 Diversity of the colibactin determinant

To find out whether the prevalence of the colibactin gene cluster is restricted to specific phylogenetic lineages of *E. coli* and *Klebsiella* spp., the sequence types of the corresponding *E. coli* and *Klebsiella* spp. isolates were further analyzed. As shown in Fig. S1, the *clb* gene cluster was enriched in a small subset of *E. coli* STs (twelve out of 11,537 STs, as of 30.10.2020), *K. aerogenes* STs (two out of 214 STs, as of 30.10.2020), and *K. pneumoniae* STs (six out of 5,237 STs, as of 30.10.2020), respectively. In these twelve *E. coli* STs, between 58% and 94% of the allocated isolates carry the *clb* determinant. A high percentage (ca. 96%) of the *K. aerogenes* ST4 and ST93 included in our study harbored the colibactin genes. In the tested *K. pneumoniae* strains, all ST3 isolates were *clb*-positive, more than 75% of the analyzed ST23 and ST234 isolates carried the colibactin gene cluster, whereas this was only the case for a significantly lower percentage of the *K. pneumoniae* isolates allocated to ST11, ST258, and ST48. Table S2 (Supplemental Material) contains a complete list of STs to which colibactin-positive *E. coli* and *Klebsiella* isolates have been assigned.

The nucleotide sequences of the *clb* gene cluster extracted from the 2,169 strains (Supplemental Material, Tab. S1) were used to generate a recombination-free phylogeny of the colibactin determinant as shown in Fig. 2A (Fig. S2 and S3 for branch support values and strain labels/assembly IDs, Fig. S4 indicates predicted recombination events in the *clb* gene cluster). *Serratia marcescens* strain MSU97 isolated from a plant source, *Erwinia oleae* strain DAPP-PG531 isolated from olive tree knot, *Klebsiella michiganensis* strain NCTC10261 of an unknown source, and the *E. coli* phylogroup E strain 14696-7 isolated from the pericardial sac of a white-tailed deer (*Odocoileus virginianus*) harbor the most genetically distant variants of the colibactin determinant (Fig. 2A). Within the *Enterobacteriaceae*, a large group of *clb* gene clusters can be defined, which is dominated by two highly conserved clades present in *E. coli* phylogenetic lineage B2 and different *Klebsiella* spp. isolates, respectively. The colibactin determinants detected in *E. cloacae* and *E. hormaechei* belong to the *Klebsiella* clades of *clb* loci, whereas the *clb* gene clusters found in *C. koseri* and in an unspecified *Salmonella* isolate represent an independent clade, i.e. *clb6* (Fig. 3). In a few other *E. coli* and *Klebsiella* spp. genomes, the *clb* determinant can be distinguished from the two major conserved clades of colibactin determinants observed in *Klebsiella* or *E. coli*. An even more divergent group comprises the *clb* gene clusters of mainly *E. coli* phylogroup A, B1, and a few B2 isolates, but also of some K. pneumoniae strains (Fig. 3, belonging to *clb* clades *clb1* and *clb2*).

**Figure 2.**
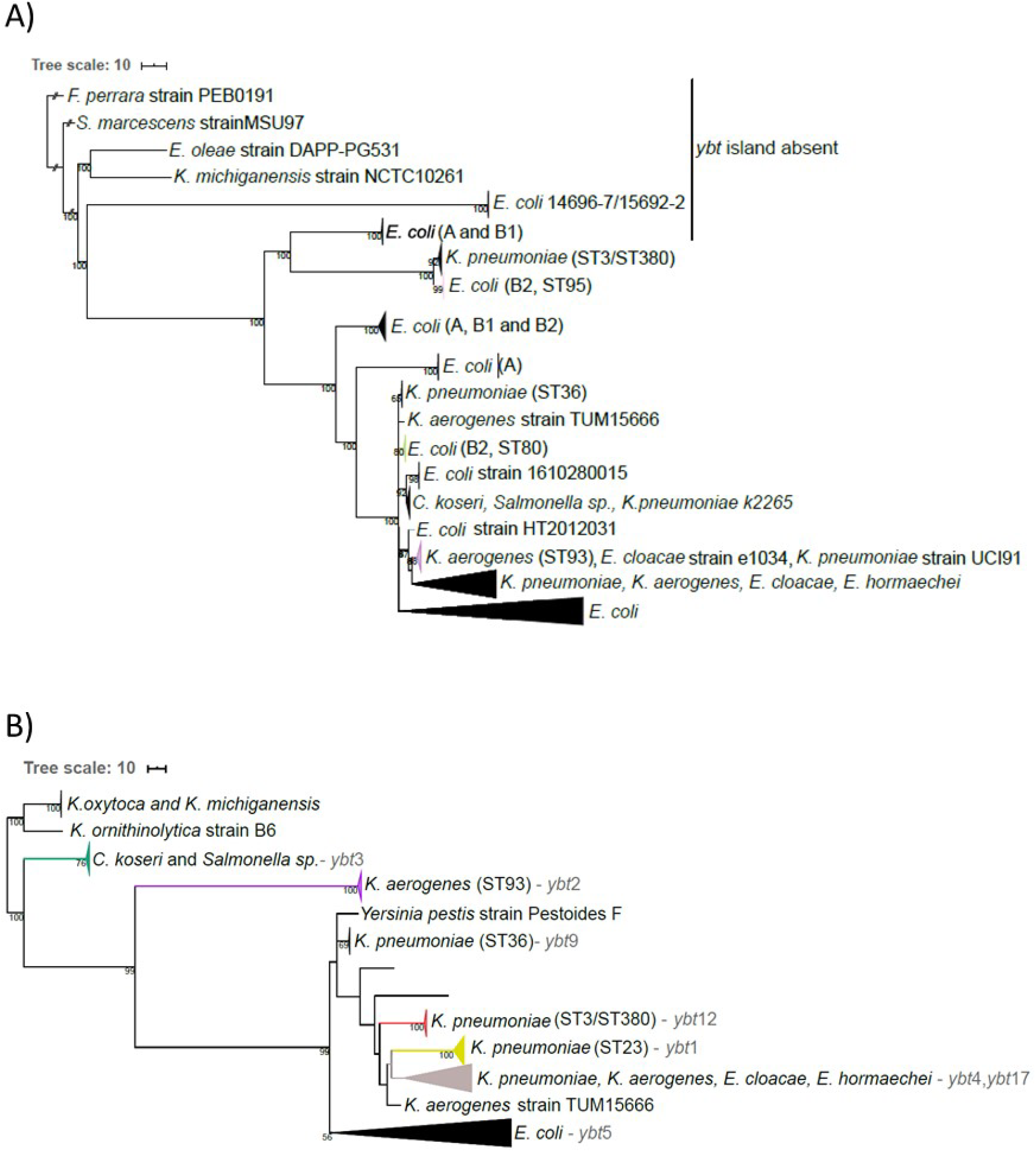
Maximum-likelihood based phylogenetic analysis of the colibactin and yersiniabactin determinants. (A) Phylogenetic tree of the colibactin gene cluster (collapsed), B) phylogenetic tree of the corresponding *ybt* determinants (collapsed) using the genetically distant *K. michiganensis* strains as outgroup (Lam *et al.*, 2018). Additionally, the yersiniabactin sequence type (YbST) as defined by Lam and colleagues (Lam et al., 2018) associated with individual bacterial clades are indicated. The branch colors in both trees depict the prominent bacterial sequence type of the clade.

**Figure 3.**
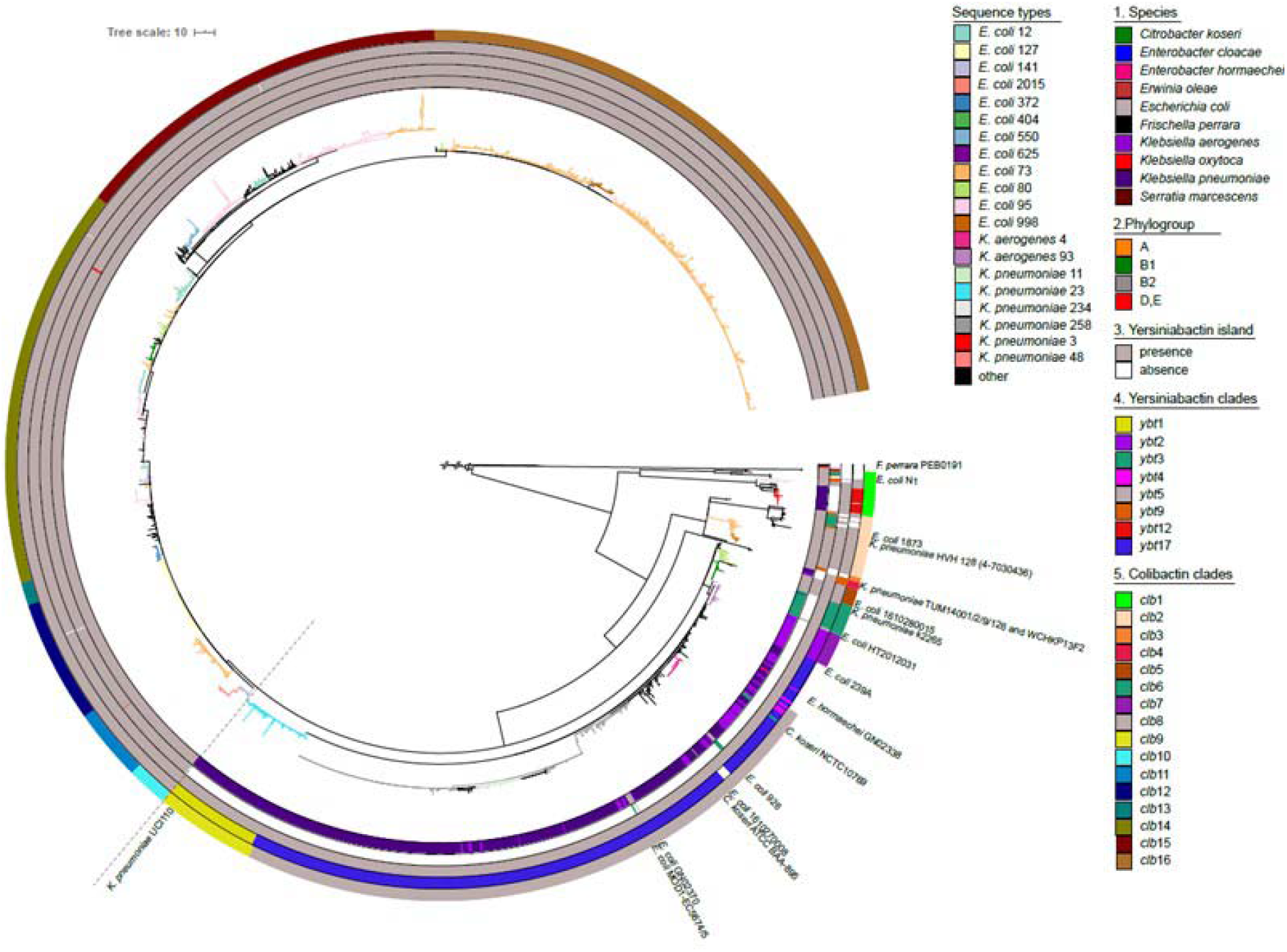
Maximum-likelihood based phylogeny of the colibactin gene cluster detected in 2169 enterobacterial genomes. Every leaf represents a single sequence variant of the *clb* gene cluster, which can be allocated to different lineages and clades. From innermost to outermost, the 1^st^ circle indicates the species harboring the *clb* determinant; the 2^nd^ circle shows the *E. coli* phylogroup, the 3^rd^ circle shows the presence/absence of the *ybt* operon; the 4^th^ circle shows the yersiniabactin sequence types (YbST) of the *ybt* determinant (from Fig. 1B) that correspond to the pks island lineage present in the individual genome. The 5^th^ circle shows the different colibactin sequence types (CbST) of the *clb* gene cluster. The branch colors in the center of the tree depict the prominent bacterial sequence types (Fig. 1). The large conserved *E. coli* phylogroup B2 clade is separated from the large *Klebsiella* clade with a faint broken line.

Although the *clb* gene clusters found in the large clade of *E. coli* B2 strains are highly conserved, individual ST-specific lineages, such as *clb*10 (from *E. coli* ST141 and ST2015 strains) and *clb*12 (from *E. coli* ST121 strains) can still be described within this clade. Beyond that, we also observed multiple lineages per sequence type, such as *clb*2, *clb*11 and *clb*16 found in *E. coli* strains of ST73 (Fig. 3). Three lineages of the *clb* locus were predominantly detectable in *Klebsiella pneumoniae* strains. They belong to the most distant *Klebsiella* ST3/ST380 clade (*clb*1), the remaining large and diverse ST258/ST11 clade (*clb*8), and finally the hypervirulent ST23 clade (*clb*9) (Fig. 3). A phylogeny of the colibactin gene cluster inferred from concatenated amino acid sequences of the 17 *clb* genes (Fig. S8) was very similar to the aforementioned recombination-free nucleotide-based phylogeny (Fig. 3).

### 5.3 Diversity of the yersiniabactin determinant in colibactin-positive bacteria

The majority (>98%) of the *clb*-positive enterobacterial strains also harbored the yersiniabactin genes (*ybt*) (Fig. 3, 3^rd^ circle). The *E. coli* strains of phylogroup A, B1, and E as well as the tested *K. michiganensis* and *E. oleae* strains carrying the most genetically distant lineages of the colibactin gene cluster, together with the *F. perrara* strains used as an outgroup, are ybt-negative (Fig. 2B). There were no strains that carried multiple copies of the colibactin gene cluster; yet two well-separated copies of the *ybt* determinant were, however, found in *C. koseri* strains ATCC BAA-895 and 0123A_53_520. It should be noted that the latter strain is derived from a metagenome (Fig. S6). The phylogenetic analysis indicated that all *ybt* operons from *E. coli* clustered together (Fig. 2B). Alike the *clb* gene cluster, also the *ybt* locus of the *E. coli* phylogroup B2 strains was highly conserved. In contrast, the sequence comparison of the *ybt* determinants of *Klebsiella* spp. resulted in different lineages, which correlate with lineages *ybt*1, 12, and 17 (ICE*Kp10*), *ybt9* (ICE*Kp3*), and *ybt*4 (originating from a plasmid) previously described by Lam and colleagues [38].

### 5.4 Congruent phylogeny of the colibactin and yersiniabactin determinants

The strong coexistence of the colibactin and yersiniabactin determinants on the one hand and the description of different evolutionary lines of *clb* and *ybt* determinants in different enterobacterial species on the other hand led us to analyze whether both gene clusters can predominantly be transferred individually or together. Our results indicate that the clades of the evolutionary lineages of the *clb* and ybt loci are chiefly species/genus-specific. The phylogeny of *clb* and *ybt* determinants is largely congruent, with the *ybt* gene cluster being, even more, species/genus-specific than that of the *clb* gene cluster. However, in some strains we observed evidence of interspecies transfer of these genes: The *clb* and corresponding *ybt* determinants of the *C. koseri* isolates NCTC10769, ATCC BAA-895 and *E. coli* strains 239A, 926, GN02370, MOD1-EC5674/5 were allocated to the large *Klebsiella*-dominated lineage clb8 (Fig. 3). Additionally, the *clb* gene cluster of *K. pneumoniae* strain k2265 was found within lineage clb6, which predominantly represents *C. koseri* isolates. However, the ybt determinant of *K. pneumoniae* strain k2265 belonged to the ybt12 lineage represented by *K. oxytoca* isolates. Similarly, the *ybt* determinant of the aforementioned strains *E. coli* strains GN02370 and *C. koseri* ATCC BAA-895 belonged to lineage *ybt*4 (plasmid originating *ybt* loci) instead of *ybt*17. Regardless of the aforementioned exceptions, clades of the *clb* gene cluster usually correlated with the corresponding clade of *ybt* genes (Fig. 3, 3^rd^ and 4^th^ circle).

### 5.5 Genetic structure of the MGEs harboring the colibactin determinant

To further investigate whether the colibactin and yersiniabactin determinants are jointly distributed by horizontal gene transfer and to obtain clues to the underlying mechanism, we compared the chromosomal context of the two polyketide biosynthesis gene clusters and the genetic structure of associated mobile genetic elements (MGEs). We observed species-specific structural differences of the chromosomal regions harboring the *clb*, and *ybt* gene clusters (Fig. 4). In *E. coli* phylogroup B2 strains, the *clb*, and *ybt* determinants are present as part of two individual pathogenicity islands (PAIs) with their cognate integrase and different tRNA genes (class I of *clb*-harboring MGE). Both PAIs are located neighboring each other in the chromosome. Within class Ia, the two PAIs are separated by a type 4 secretion system (T4SS)-encoding operon (*virB*) and a region that includes two conserved gene sets (Set 1 and Set 2), different tRNA(*Asn*) loci and an integrase gene (see Table S3 for genes present in the different conserved gene sets). This region is shown to have been diminished in class Ib structural variants, where only gene *virB1* of the T4SS determinant is left alongside the conserved region. In the predominant *E. coli* structural variant, class Ic, the complete T4SS operon (including *virB1*) has been lost together with gene set 1, the integrase gene was exchanged and a DNA stretch comprising the genes *yeeO*, tRNA(*Asn*), *cbl* and *gltC* was inverted (shown in red, Fig. 4). The region separating the *pks* island and the HPI was reduced from a 40-kb (in class Ia) to a 15-kb stretch in class Ic. In contrast, within all *Klebsiella* strains, the *clb* and *ybt* gene clusters are part of an ICE, and are separated by a T4SS-encoding operon (*virB*) and the *hha* gene coding for the hemolysin expression-modulating protein. Downstream of the *ybt* gene cluster an integrase gene is located followed by a set of genes involved in Fe/Mn/Zn metabolism (structural class II of *clb*- and *ybt*-harboring chromosomal regions) (Fig. 4). Interestingly, one type of such ICEs, is located next to genes necessary for microcin E492 biosynthesis (class IIc, Fig. 4). *Enterobacter hormaechei* strains harbor a structurally similar ICE to that of *Klebsiella* strains. In most *C. koseri* strains, however, the two polyketide determinants are separated by a large 250-kb chromosomal region. The T4SS-related genes are closely positioned to the *clb* genes while the gene set involved in Fe/Mn/Zn metabolism is located downstream of the *ybt* determinant. Only a minor fraction of enterobacterial isolates analyzed displayed some variation regarding gene content and synteny of these three main classes of colibactin and yersiniabactin-encoding chromosomal regions. The structure of *clb* and *ybt* regions that do not conform to these major classes are as shown in Figure S5. Instead of class I, several *E. coli* strains carried class II like chromosomal regions where the T4SS and the Fe/Mn/Zn metabolism genes were present. In *E. coli* strain HVH128 none of the three main classes colibactin and yersiniabactin-encoding regions could be identified. Although both polyketide determinants are co-localized with one integrase gene each, they are widely separated on this strain’s chromosome. *K. pneumoniae* strains TUM14001, TUM14002, TUM14009, TUM14126, and WCHKP13F2 harbored two T4SS-encoding gene clusters in close proximity of the *clb* and *ybt* gene clusters and lacked the Fe/Mn/Zn genes, whereas *K. pneumoniae* strain UCI110 was also missing the Fe/Mn/Zn metabolism-related genes. In contrast to the other *C. koseri* isolates, we detected a class II- instead of a class III-type *clb*-*ybt* region in *C. koseri* isolate BAA-895.

**Figure 4.**
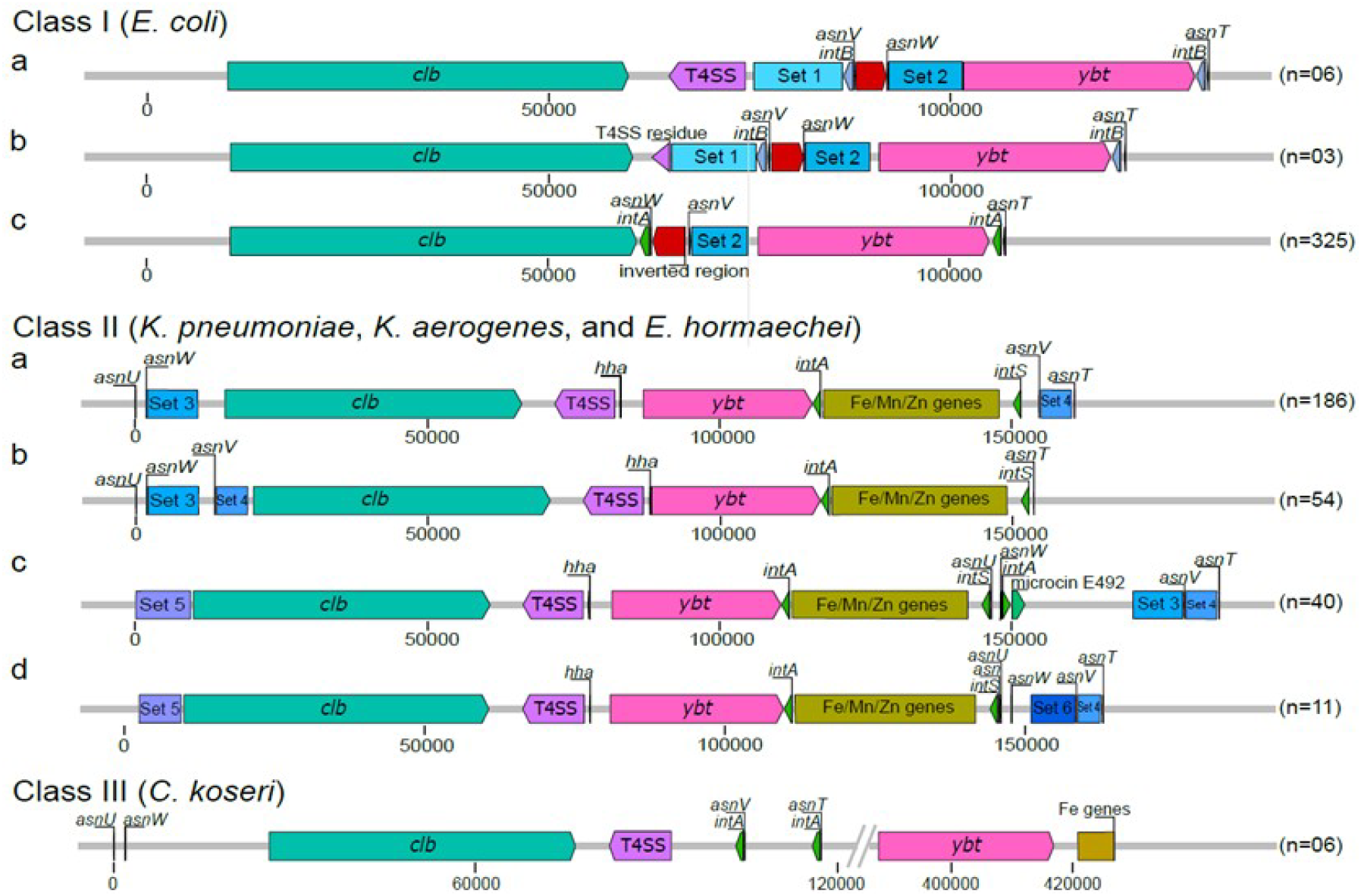
Structural variation of the colibactin and yersiniabactin-encoding chromosomal region in *E. coli*, *E. hormaechei*, *K. pneumoniae*, *K. aerogenes* and *C. koseri*. The different genetic structures and chromosomal insertion sites of the colibactin and/or yersiniabactin determinants found within the three main structural classes are shown. The *clb* gene cluster (teal green), T4SS module (purple), *ybt* gene cluster (pink), integrase genes (green), the conserved sets of genes (Table S2) that are present up/downstream the two polyketide determinants, classed into sets (blue boxes) and the Fe/Mn/Zn module (yellow) are shown. The number of genomes included in the tested set of genomes that harbour the different structural variants is indicated in brackets. The colibactin-yersiniabactin chromosomal regions that do not conform to these major structures are as shown in Figure S4.

### 5.6 Organization of the colibactin gene cluster

The *clb* gene cluster is composed of 19 genes, which are required for the regulation, biosynthesis, and transport of colibactin. The origin of this gene cluster in unclear. We, therefore, compared the structure of the gene clusters representing the homologous *clb* locus found in *F. perrara* and the phylogenetically most distant and potentially older *clb* determinants relative to the *E. coli* B2 type of *clb* determinant (Fig. 2A), which are present in *S. marcescens*, *E. oleae*, *K. michiganensis*, and *E. coli* phylogroup E strain 14696-7(Fig. 5). The various *clb* determinants correspond in terms of the structure of the genes coding for the biosynthesis machinery, transport, and resistance to colibactin (clbB-clbS) and resemble the structure of the well-described *clb* locus in *K. pneumoniae/K. aerogenes/E. coli* B2 strains. The individual *clb* determinants differ more clearly in the presence and localization of the genes involved in the regulation and activation of colibactin biosynthesis (*clbR* and *clbA*). These two genes are absent in the homologous gene cluster found in *F. perrara* and in the *clb* determinant in *S. marcescens*. However, in *F. perrara*, a phosphopantetheinyl transferase coding for a homolog of ClbA (43% amino acid similarity) and a radical S’-adenosylmethionine (SAM) enzyme-encoding gene are found directly downstream of clbS. In *S. marcescens* a helix-turn-helix (HTH)-type regulatory protein homologous to clbR is encoded by a gene located upstream of clbB (78% amino acid similarity). In *E. oleae* and *K. michiganensis* a SAM enzyme-encoding gene is present directly downstream of *clbA*. Although the colibactin gene clusters in *E. oleae* and *K. michiganensis* have a high nucleotide sequence similarity of ca. 99.77%, the predicted coding regions of *clbC/D* and *clbH/I/J/K/L/M/N* are noticeably different due to multiple frameshift deletions in *K. michiganensis*. The structure of the *clb* locus found in phylogroup E *E. coli* strain 14696-7 already corresponds to the structure of *E. coli* strains of phylogroup B2 (Fig. 1 and Fig. 5), yet this gene cluster shows the least sequence similarity of all tested *clb* gene clusters in non-B2 *E. coli* isolates (Fig. 2A) to the determinant occurring in *E. coli* strains of phylogroup B2.

**Figure 5.**
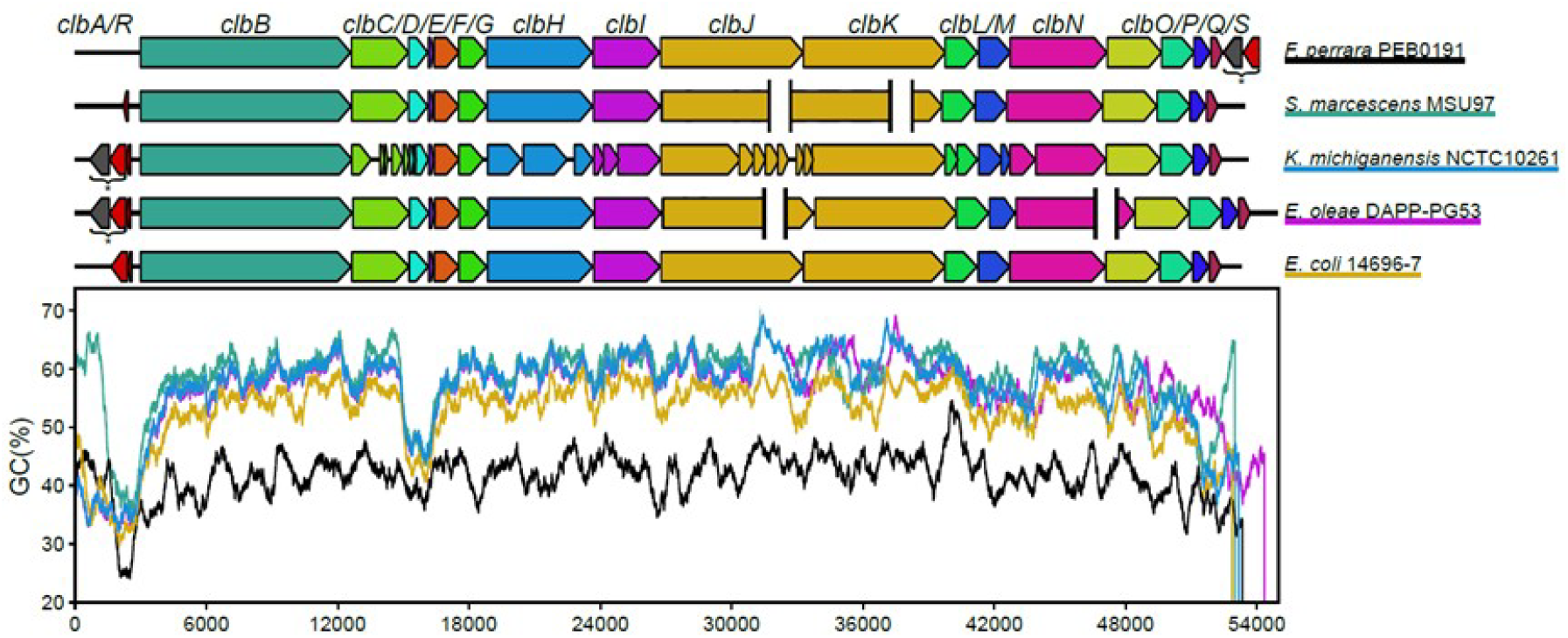
The structural organization and GC profiles of the *clb* determinants in the five most genetically distant bacterial strains (according to Fig. 2A). The genes that make up the homologous *pks* gene cluster found in *F. perrara* and the most distant *clb* determinants present in *S. marcescens*, *E. oleae*, *K. michiganensis* and *E. coli* (phylogroup E) strain 14696-7 are depicted. The GC profile of the gene cluster in the different strains is shown alongside with the colors underlining the different species. SAM genes and *clbA* homologues (*) are shown downstream of the *pks* gene cluster in *F. perrara* and upstream of the *clb* determinant in *K. michiganensis* and *E. oleae*. The gaps in assembly are shown with white spaces.

Looking at the G+C plot of the colibactin gene clusters, it is obvious that all investigated enterobacterial *clb* gene clusters show a very similar G+C plot, which has a significantly higher average G+C content and differs significantly from that of the *clb* homologous gene cluster in *F. perrara* (Fig. 5). The G+C content profile of these gene clusters indicates that there are two regions of low G+C content in the enterobacterial *clb* determinants: the region including *clbA* and *clbR* (at position ca. 1,500 bp – 3,000 bp of the colibactin gene cluster) and the region spanning *clbD* and *clbE* (at position ca. 15,000 bp – 16,500 bp of the *clb* gene cluster). The G+C content drop in the region including *clbA* and *clbR* (at position ca. 1,500 bp - 3,000 bp of the colibactin gene cluster) is associated with a predicted recombination site, which is located upstream of or interrupting *clbB* (Fig. S4).

The comparison of structural features of the *clb* gene cluster also included the VNTR region located upstream of *clbR* in the *clbR-clbB* intergenic region. The size of the VNTR region has been described to range from 2 to 20 (Putze *et al.*, 2009). The VNTR copy number distribution in ca. 1,300 *clb*-positive genomes demonstrated that there is a preference for VNTR regions ranging from 7-10 copy numbers. Copy numbers from 18-34 were present in only a few strains (Figure S6). Species and/or ST-specific copy number variation was not observed.

Comparative genomic analysis of multiple colibactin-encoding determinants based on (draft) genome sequences led to the observation that the homologous genes *clbJ* and *clbK* are prone to fusion/deletion (Lam *et al.*, 2018). We also observed that in several assemblies of the *clb* gene cluster 625 bp from the 3’ end of *clbJ* and 3,540 bp from the 5’ end of *clbK* including the 11 bp intergenic region are missing, for a total of 4,174 bp (Fig. S5). The assembly of our internally generated genome sequences produced by short read (Illumina) sequencing showed this *clbJ/K* fusion/deletion. However, since assemblies of sequence data of the same strains generated by a long-read sequencing technology (PacBio), where the long reads covered both genes, had both *clbJ* and *clbK* completely present, we assume that the *clbJ/K* fusions described are artificial and result from erroneous assemblies of short-read sequencing data.

### 5.7 Quantification of colibactin synthesis in selected strains

To investigate a possible correlation between the genetic structure of the *clb* determinant or the genetic background of the corresponding host strain with colibactin expression, we quantified N-myristoyl-D-asparagine levels produced *in vitro* by selected clb-positive *E. coli*, *Klebsiella* spp. and *C. koseri* strains covering the diversity of *clb* determinants in these species (Fig. 6). Based on the detected relative amount of N-myristoyl-D-asparagine produced, the investigated isolates can be roughly divided into two groups: One group included most of the measured *E. coli* strains that produced only very low relative amounts of N-myristoyl-D-asparagine. In contrast, the tested *C. koseri*, *K. aerogenes*, and *K. pneumoniae* isolates and the *E. coli* isolates CFT073 and N1 showed a 3 to 70-fold higher N-myristoyl-D-asparagine production. Also within the *C. koseri*, *K. aerogenes*, and *K. pneumoniae* isolates, we found differences in the relative N-myristoyl-D-asparagine levels. However, these differences were not as strong as among the eight *E. coli* isolates studied. The observed relative N-myristoyl-D-asparagine levels do not indicate phylogroup, ST, or species-specific differences in colibactin production. For example, the *E. coli* strain 1873, although the *clb* gene cluster present in this strain is phylogenetically more closely related to that of *E. coli* strain N1, shows a significantly weaker N-myristoyl-D-asparagine production than *E. coli* N1. Similarly, it should be noted that *C. koseri* MFP3 produces less N-myristoyl-D-asparagine than other closely related *C. koseri*.

**Figure 6.**
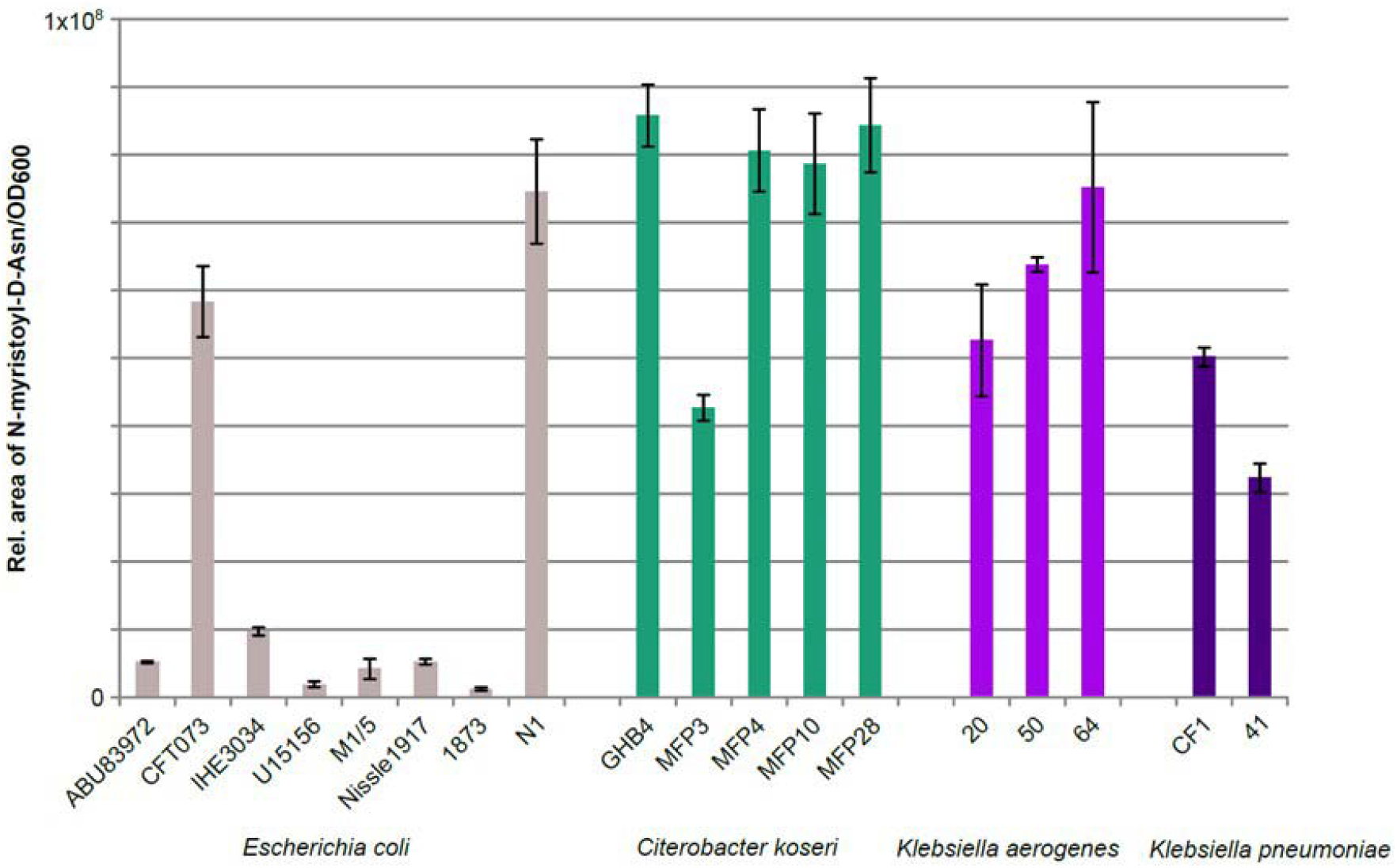
Comparison of colibactin production of different strains accessed by quantification of the precolibactin cleavage product N-Asn-D-myristol. This assay enabled us to compare the ability of different strains and different species to produce colibactin under controlled conditions *in vitro*. Measurements were conducted based on three biological replicates, means with standard deviations are as shown.

## 6 Discussion

### *Prevalence and mobility of the colibactin gene cluster in* Enterobacteriaceae

The ability of MGEs to be exchanged between and within species plays a major role in the extent and speed of microbial evolution. Because MGEs are known to emerge and evolve separately from the host, it is important to explain the development and diversity of MGEs independently. In our study, we investigated on the one hand the nucleotide sequence variability of the colibactin gene cluster with the associated yersiniabactin determinants, and on the other hand the structural diversity of the MGEs hosting the colibactin and yersiniabactin determinants responsible for their horizontal distribution. Both polyketide determinants together could be detected in certain members of the *Enterobateriaceae*, mainly in *E. coli*, *K. pneumoniae*, *K. aerogenes*, and *C. koseri*, but also in *Enterobacter cloacae*, *Enterobacter hormachei*, and possibly in uncharacterized *Salmonella* sp. isolates. The colibactin, but not the yersiniabactin determinants could also be detected in *K. michiganensis*, *E. oleae*, and *S. marcescens* and a few phylogroup A and B1 *E. coli* isolates (Fig. 2A). Interestingly, the colibactin genes have not yet been described in not only archaeal genomes but also *S. enterica* and *S. bongori* genomes, although we have screened more than 112,000 *Salmonella* spp. genomes. Overall, it is remarkable that the *clb* gene cluster was only found in extraintestinal clinical or in fecal isolates of healthy hosts, but not in enterobacterial diarrhoeal pathogens such as Yersinia spp., *Salmonella* spp., *Shigella* spp. as well as the various intestinal pathogenic *E. coli* pathotypes. This observation is consistent with previously published data [2, 50]. In this context, it would be interesting to study in the future whether and in which way colibactin expression supports extraintestinal pathogenicity or intestinal persistence and colonization, but is detrimental to the pathogenesis of diarrhoeal pathogens. Possibly, the genetic background also plays an important role in the horizontal distribution and establishment of MGEs carrying both polyketide determinants. The fact that the colibactin determinant has so far been preferentially distributed in only some, often highly virulent STs in *E. coli* and *K. pneumoniae* or *K. aerogenes* and not more broadly within the respective species (Fig. 2) [38, 39], could also indicate that the transmission, uptake or chromosomal integration of these MGEs is restricted. It is interesting to note that the *clb* gene cluster as a whole is highly conserved and usually characteristic of the respective species or genus (Fig. 3). Nevertheless, ST-specific variants have been found within a species, e.g. in *K. pneumoniae* ST3 and ST23. Several groups of sequence variants of the *clb* gene cluster can also be found within one ST, such as in the *E. coli* ST73 and ST95 (Fig. 3). These results show that, on the one hand, intraspecies transfer of the colibactin determinant can happen, but on the other hand, certain adaptations of the *clb* genes to a specific genetic background can also occur at the nucleotide level. In addition, examples of interspecies transfer of the *clb* and *ybt* genes can be seen between *Klebsiella* spp. and *E. hormachei* strains. Similarly and in contrast to the majority of *E. coli* isolates, we found *clb* and *ybt* gene clusters in some *E. coli* isolates, which are assigned to the large clade of *Klebsiella/Enterobacter/Citrobacter*-specific *clb* and *ybt* variants (Fig. 2A and 3). Apart from the interspecies transfer of the entire *clb*- and *ybt*-containing MGE, we also identified an example that shows that the two polyketide gene clusters can also be exchanged independently, as in the case of the *clb* gene cluster of *K. pneumoniae* strain k2265, which belongs to colibactin clade *clb*6 (predominantly *C. koseri* lineage of *clb* loci), whereas the *ybt* determinant in this strain is assigned to the yersiniabactin clade *ybt*12 instead of *ybt*3, which is usually associated with *C. koseri* strains carrying clade clb6 (Fig. 3).

### Structural diversity of the colibactin-yersiniabactin region

The structural analysis of the genomic region comprising the *clb* and *ybt* determinants in their chromosomal sequence context is an important aspect to understand the evolution of these polyketide determinants and their origin. In principle, three structural constellations (classes I to III) can be described, in which the *clb* and *ybt* gene clusters are present (Fig. 4). Class I depicts the *clb* and *ybt* gene clusters found in the majority of *E. coli* isolates, each associated with a tRNA(Asn) and an integrase gene. In class Ia, the *clb* genes are chromosomally inserted at the tRNA locus asnV. Our analyses suggest that in class I, the type 4 secretion system gene cluster (“mobilization module”) and conserved neighboring genes (set 1) have been lost in a stepwise process, from class Ia to Ic. A further structural modification in this region is represented by the inversion of the *yeeO-tRNA(Asn)-cbl-gltC* gene set (Fig. 4, red arrow), as a result of which in class Ic, the tRNA gene *asnW* is located closest to the *clb* genes. Taking into account that the *E. coli* strains with class Ia and Ib structures are found in the potentially earlier phylogenetic clades, we hypothesize that the MGE harboring the *clb* genes was introduced into the *E. coli* chromosome separately from that carrying the *ybt* determinant. Both MGEs were then progressively modified as described above. We do not yet have an explanation why the resulting class Ic colibactin-yersiniabactin region, which has been described as two pathogenicity islands (PAIs) comprising the colibactin and yersiniabactin determinants, respectively [1, 2], is only found in phylogroup B2 strains and only there has it become so successful.

Class II includes different variants of an ICE, in which the two polyketide determinants are present in association with a type 4 secretion system-encoding “mobilization module” and a “module” consisting of genes that contribute to Fe/Mn/Zn metabolism (Fig. 4, Supplementary Table S3). This type of ICE was found in *Klebsiella* spp. and *E. hormachei* and in a few cases in *E. coli* and *C. koseri*. Unlike in class I and III, the ICE in class II does not only have different tRNA(Asn) loci serving as chromosomal insertion sites, but also lacks tRNA(*Asn*) and integrase genes in between the *clb* and *ybt* genes. In a population-wide analysis of *Klebsiella* spp. strains, this ICE was designated ICEKp10 and described as being associated with different 17 combinations of *ybt* and *clb* gene lineages [38]. In contrast to class I and II colibactin-yersiniabactin regions, the *clb* and *ybt* gene clusters are located far apart on the chromosome in most of the *C. koseri* genomes studied (class III) (Fig. 4).

The existence of an ICE that unites the *clb* and *ybt* gene clusters is the easiest way to explain the co-localization and the joint transfer of the two determinants and thus the high correlation of *clb* and *ybt* phylogenetic clades (Fig. 3, 4^th^ circle) in *Klebsiella* spp. strains. Despite slight differences in the sequence context and different chromosomal insertion sites (Fig. 4), the ICEs of the four class II variants have an overall identical genetic structure (Fig. 4). The uptake of this ICE thus leads to the acquisition of both the *clb* and the *ybt* gene clusters. The presence of the Fe/Mn/Zn metabolic genes neighboring these ICE variants with an additional integrase gene indicates recombination processes that can alter the genetic structure of the ICE. The clear separation between the *clb*/T4SS module and the *ybt*-Fe/Mn/Zn metabolism module in the *C. koseri* genomes points towards rearrangement/relocation of the *ybt* region, after ICE integration into the chromosome. The fact that *C. koseri* strain ATCC BAA-895 possesses in addition to the complete ICE (class II) a second *ybt* gene cluster (99.98% nucleotide similarity to the *ybt* genes present in the ICE) that is located far away from the ICE (Fig. S6), supports the hypothesis that the individual polyketide gene clusters can also be integrated into the genome independently of each other. This state could result, for example, from the initial chromosomal integration of different ICE variants, as described by Lam and colleagues in *Klebsiella* spp. [38]. As a result of deletion events, through which individual modules are subsequently deleted from one of the two ICEs, the second copy of the *ybt* gene cluster remains in the genome as a fragment of the degenerated ICE. The presence of two non-identical T4SS modules in the *K. pneumoniae* strains TUM14001, TUM14002, TUM14009, TUM14126, and WCHKP13F2 (Fig. S6 and Fig. 3, *clb*4) associated with the *clb* or *ybt* module (Fig. S6) could be the result of such degeneration of different ICEs. In this way, our observations on the phylogeny (Fig. 2) and structure (Fig. 4) of the two polyketide determinants and their sequence context can be reconciled, which show that despite the predominant species/genus specificity, there are also sequence type-specific lineages of the *clb* genes, which do not necessarily have to match that of the associated *ybt* genes.

In this context, one could imagine that the arrangement of *clb* and *ybt* determinants in *E. coli* strains (class I) also results from independent integration events of different MGEs, which subsequently degenerated as a result of a stabilization process of these MGEs [26, 28]. This premise is supported by the absence of *ybt* genes in the clb-positive *E. coli* strains, which carry the phylogenetically most distant *clb* determinants compared to the *clb* genes of phylogroup B2 isolates (Fig. 2A), along with the presence of integrase and tRNA(Asn) genes in close proximity to both polyketide determinants (Fig. 4, class I).

### Evolution of the colibactin determinant

Homologs of the *clb* gene cluster were detected in marine alpha-proteobacteria such as various Pseudovibrio sp. (isolates AD26, FO-BEG1, POLY-S9) and *Pseudovibrio denitrificans* (isolates DSM 17465 and JCM12308) [51]. Despite the general conformity of the genetic structure of these gene clusters, their nucleotide sequence identity to the colibactin gene cluster is quite low (<26%). Therefore, it was hypothesized that these *Pseudovibrio* isolates have the potential to produce molecules related to colibactin [51]. Another homolog of the colibactin determinant with a higher (62%) amino acid sequence identity is found in *F. perrara* [24]. While the genes required for biosynthesis and transport of the polyketide are present, the genes corresponding to *clbA* and *clbR* are missing in this gene cluster (Fig. 5). The case is similar with the colibactin gene cluster in *S. marcescens*. While in *F. perrara*, a gene coding for a *clbA* homolog and a gene coding for a SAM enzyme are located immediately downstream of the *clbS* homolog, a *clbR* homolog is located upstream of *clbB* in *S. marcescens*. It is therefore conceivable that the *clbA* homolog in *F. perrara* and the *clbR* homolog in *S. marcescens* are involved in the activation or regulation of colibactin biosynthesis in these bacteria (Fig. 5). It has been described that S-adenosylmethionine (SAM) is used in NRPS modules for colibactin biosynthesis [52]. Looking at the genetic structure of the *clbB-S* homologous genes cluster in *F. perrara* and the *clb* gene cluster in *S. marcescens*, one can assume that genes involved in the regulation and activation of the biosynthetic pathway including the SAM enzyme gene as well as *clbA* and *clbR* homologs have been fused upstream to the already existing part of the island (*clbB-clbS*) to improve regulation of polyketide biosynthesis. Without having further knowledge about the origin of the colibactin biosynthetic genes themselves, the acquisition of the regulatory/activating genes is obviously among the last evolutionary steps that led to the structural organization of what we currently describe as the colibactin gene cluster. This hypothesis is supported by the abrupt decrease in G+C content and the presence of the predicted recombination (Fig. S4) directly upstream of *clbB* in many *clb* determinants. Furthermore, a module consisting of a gene for a SAM enzyme and a *clbA* homolog is not only located directly downstream from the *clbS* homolog in *F. perrara*, but also upstream from *clbR* in *K. michiganensis* and *E. oleae*, which represent evolutionarily older variants of colibactin-positive Enterobactericeae (Fig. 5).

### Expression of colibactin in different hosts

Furthermore, we investigated the question of how differently colibactin is expressed within enterobacterial genera or even within different lineages of the same species. Interestingly, we observed an often lower production level of N-myristoyl-D-asparagine in *E. coli* isolates compared to *K. aerogenes*, *K. pneumoniae*, and *C. koseri* (Fig. 6), which may be expected since *E. coli* is described as a non-optimal producer of complex secondary metabolites [53]. However, it is of interest that the amount of N-myristoyl-D-asparagine produced in *E. coli* strains CFT073 and N1 is comparable to that of other enterobacterial genera (Fig. 6). A species- or lineage-specific ability to produce N-myristoyl-D-asparagine could not be determined so far. Future studies will have to investigate which bacterial factors are important for colibactin production and how the strain-specific differences in the expression of this polyketide come about. The systematic comparison of phenotypic colibactin production with information on the genomic context, regulatory and metabolic properties of host strains, and their classification in a phylogenetic context should help us to identify bacterial factors that affect colibactin synthesis.

## 7 Conclusion

The colibactin and yersiniabactin gene clusters are highly conserved polyketide determinants within some *Enterobacteriaceae*. They usually coexist together in the genome and are also linked to each other at the biosynthetic level. With the exception of *E. coli*, the two gene clusters are part of an ICE, which allows the horizontal transfer of both secondary metabolite determinants usually within one species/genus. Bacteria of the genus *Klebsiella* played an important role in the evolution and distribution of both gene clusters. A large number of different ICEs has been described in *Klebsiella* spp., which besides several other groups of genes include the yersiniabactin determinant [38]. Recombination and rearrangements events between different ICE types may have contributed to the evolution of the ICE variants so far identified in *Klebsiella* spp. and other enterobacteria as well as to the further degeneration of such MGEs leading to the colibactin and yersiniabactin-encoding PAIs present in phylogroup B2 *E. coli* strains (Fig. 7).

**Figure 7.**
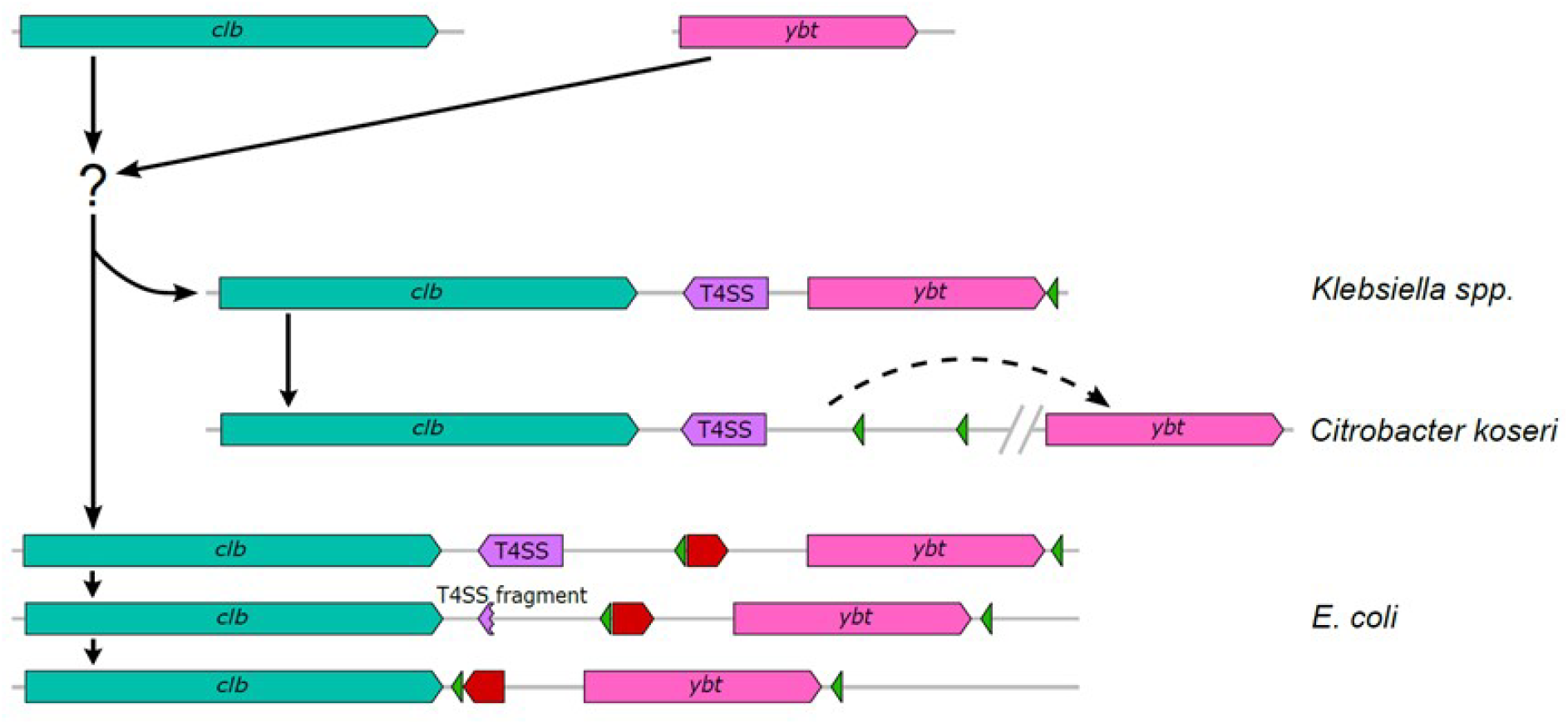
Schematic representation of the predicted evolution of the colibactin-yersiniabactin genomic region in *Enterobacteriaceae*. The different elements of this region, i.e. the *clb* determinant (teal green), T4SS module (purple), *ybt* gene cluster (pink), integrase genes (green), and an invertible subset of genes (red arrow) are shown. Based on available genome sequence data, we suggest a development from single MGEs containing the *clb* determinant and the *ybt* gene cluster, respectively, towards the structural arrangement of both polyketide determinants, which is now mainly found in enterobacterial populations. Black arrows (solid or dashed) indicate possible directions of development and DNA rearrangements. After the merge of the *clb* and *ybt* gene clusters into one MGE, represented by ICE*Kp10*, there is evidence that three different structural variants have evolved from it: In *Klebsiella* spp. strains, the ICE*Kp10* has remained intact, whereas in *C. koseri* strains, a DNA rearrangement and re-localization of the *ybt* determinant to a different chromosomal position has taken place. In *E. coli*, a gradual loss of the T4SS module and the inversion of a gene set between the two polyketide determinants led to immobilisation or stabilisation of the ICE thus resulting the two pathogenicity islands known as pks island and HPI, respectively.

**Table 1.** Bacterial species tested positive for the presence of the colibactin determinant.

|  | No. of strains screened | *clb*-positive strains |
| --- | --- | --- |
| *Escherichia coli* | 19,183 | 1,462 |
| *Klebsiella pneumoniae* | 8,040 | 572 |
| *Klebsiella aerogenes* | 225 | 101 |
| *Citrobacter koseri* | 48 | 27 |
| *Enterobacter hormaechei* | 711 | 2 |
| *Enterobacter cloacae* | 610 | 2 |
| *Serratia marcescens* | 515 | 1 |
| *Klebsiella michiganensis* | 94 | 1 |
| *Salmonella* sp. | 8 | 1* |
| *Erwinia oleae* | 1 | 1 |
| *Frischella perrara* | 3 | 3 |

\*unverified source organism (excluded from Refseq)

The phylogeny of the *clb* determinants does not determine the level of phenotypic colibactin production. The underlying bacterial factors responsible for the colibactin production efficiency of individual strains need to be identified in future work.

Our investigations provide deeper insights into the evolution of the colibactin gene cluster in *Enterobacteriaceae*. Based on our findings, we can extend the current explanation for the co-existence and genetic co-localization of both gene clusters. The combination of a PPTase-encoding gene (*clbA*) with the *clbB-S* biosynthetic gene cluster during the evolution of the *clb* determinant not only enabled the efficient activation of the colibactin biosynthesis machinery, but also linked the colibactin and yersiniabactin determinants, which are functionally connected by the activity of PPTase ClbA. This enables the bacteria to synthesize both functionally different secondary metabolites, which leads to a stabilization of the co-existence and co-localization of the two gene clusters in the genome. Our data underpin the importance of mobile genetic elements, especially of ICEs, for genomic diversity and variability in enterobacteria as well as for the evolution of more complex bacterial phenotypes, such as the combined expression of the secondary metabolites colibactin and yersiniabactin.

## Author statements

### Authors and contributors

H.W., A.W. and U.D. conceptualized the project. H.W., A.W., D.S. ran the analyses. R.M., M.S., and E.O. contributed reagents and new tools. H.W., A.W., and U.D. analysed the data. H.W., A.W. and U.D. wrote the manuscript. H.W., A.W., M.S., R.B., E.O., R.M. and U.D. edited and revised the manuscript. All authors read, commented on and approved the final manuscript.

## Acknowledgments

Data reported in this study appear in part in the PhD theses of A. Wallenstein and H. Wami. We thank M.K. Mammel (United States Food and Drug Administration) for providing *E. coli* strain N1. We thank K. Tegelkamp and O. Mantel (Münster) for excellent technical support. We gratefully acknowledge CPU time on the high performance cluster PALMA@WWUMünster.

## Supplemental Material

includes Supplementary Tables S1 (*clb*-positive bacterial strains), S2 (clb-positive STs in *E. coli* and *Klebsiella*), S3 (conserved gene sets associated with the *clb* and *ybt* determinants), S4 (list of prokaryotic species included into the *clb* screen), S5 (list of ClbSTs), and S6 (list of YbSTs)

## References

1. Nougayrède JP, Homburg S, Taieb F, Boury M, Brzuszkiewicz E et al. Escherichia coli induces DNA double-strand breaks in eukaryotic cells. Science 2006;313(5788):848–851.

2. Putze J, Hennequin C, Nougayrède JP, Zhang W, Homburg S et al. Genetic structure and distribution of the colibactin genomic island among members of the family Enterobacteriaceae. Infect Immun 2009;77(11):4696–4703.

3. Xue M, Kim CS, Healy AR, Wernke KM, Wang Z et al. Structure elucidation of colibactin and its DNA cross-links. Science, 2019;365(6457).

4. Homburg S, Oswald E, Hacker J, Dobrindt U. Expression analysis of the colibactin gene cluster coding for a novel polyketide in Escherichia coli. FEMS Microbiol Lett 2007;275(2):255–262.

5. Martin P, Marcq I, Magistro G, Penary M, Garcie C et al. Interplay between siderophores and colibactin genotoxin biosynthetic pathways in Escherichia coli. PLoS Pathog 2013;9(7):e1003437.

6. Wallenstein A, Rehm N, Brinkmann M, Selle M, Bossuet-Greif N et al. ClbR Is the Key Transcriptional Activator of Colibactin Gene Expression in Escherichia coli. mSphere 2020;5(4):e00591–00520.

7. Healy AR, Wernke KM, Kim CS, Lees NR, Crawford JM et al. Synthesis and reactivity of precolibactin 886. Nat Chem 2019;11(10):890–898.

8. Li ZR, Li J, Cai W, Lai JYH, McKinnie SMK et al. Macrocyclic colibactin induces DNA double-strand breaks via copper-mediated oxidative cleavage. Nat Chem 2019;11(10):880–889.

9. Li ZR, Li J, Gu JP, Lai JY, Duggan BM et al. Divergent biosynthesis yields a cytotoxic aminomalonate-containing precolibactin. Nat Chem Biol 2016;12(10):773–775.

10. Bossuet-Greif N, Vignard J, Taieb F, Mirey G, Dubois D et al. The colibactin genotoxin generates DNA interstrand cross-links in infected cells. mBio 2018;9(2):e02393–02317.

11. Cuevas-Ramos G, Petit CR, Marcq I, Boury M, Oswald E et al. Escherichia coli induces DNA damage in vivo and triggers genomic instability in mammalian cells. Proc Natl Acad Sci USA 2010;107(25):11537–11542.

12. Buc E, Dubois D, Sauvanet P, Raisch J, Delmas J et al. High prevalence of mucosa-associated *E. coli* producing cyclomodulin and genotoxin in colon cancer. PLoS One, 2013;8(2):e56964.

13. Cougnoux A, Dalmasso G, Martinez R, Buc E, Delmas J et al. Bacterial genotoxin colibactin promotes colon tumour growth by inducing a senescence-associated secretory phenotype. Gut 2014;63(12):1932–1942.

14. Dalmasso G, Cougnoux A, Delmas J, Darfeuille-Michaud A, Bonnet R. The bacterial genotoxin colibactin promotes colon tumor growth by modifying the tumor microenvironment. Gut Microbes 2014;5(5):675–680.

15. Dziubanska-Kusibab PJ, Berger H, Battistini F, Bouwman BAM, Iftekhar A et al. Colibactin DNA-damage signature indicates mutational impact in colorectal cancer. Nat Med 2020;26(7):1063–1069.

16. Marcq I, Martin P, Payros D, Cuevas-Ramos G, Boury M et al. The genotoxin colibactin exacerbates lymphopenia and decreases survival rate in mice infected with septicemic Escherichia coli. J Infect Dis, 2014;210(2):285–294.

17. Pleguezuelos-Manzano C, Puschhof J, Rosendahl Huber A, van Hoeck A, Wood HM et al. Mutational signature in colorectal cancer caused by genotoxic pks(+) *E. coli*. Nature 2020;580(7802):269–273.

18. Faïs T, Delmas J, Barnich N, Bonnet R, Dalmasso G. Colibactin: More Than a New Bacterial Toxin. Toxins 2018;10(4).

19. Wassenaar TM. *E. coli* and colorectal cancer: a complex relationship that deserves a critical mindset. Crit Rev Microbiol, Review 2018;44(5):619–632.

20. Wilson MR, Jiang Y, Villalta PW, Stornetta A, Boudreau PD et al. The human gut bacterial genotoxin colibactin alkylates DNA. Science, 2019;363(6428).

21. Brzuszkiewicz E, Brüggemann H, Liesegang H, Emmerth M, Olschlager T et al. How to become a uropathogen: comparative genomic analysis of extraintestinal pathogenic Escherichia coli strains. Proc Natl Acad Sci U S A 2006;103(34):12879–12884.

22. Bellanger X, Payot S, Leblond-Bourget N, Guedon G. Conjugative and mobilizable genomic islands in bacteria: evolution and diversity. FEMS Microbiol Rev 2014;38(4):720–760.

23. Schubert S, Rakin A, Karch H, Carniel E, Heesemann J. Prevalence of the “high-pathogenicity island” of Yersinia species among Escherichia coli strains that are pathogenic to humans. Infect Immun 1998;66(2):480–485.

24. Engel P, Vizcaino MI, Crawford JM. Gut symbionts from distinct hosts exhibit genotoxic activity via divergent colibactin biosynthesis pathways. Appl Environ Microbiol 2015;81(4):1502–1512.

25. Moretti C, Hosni T, Vandemeulebroecke K, Brady C, De Vos P et al. Erwinia oleae sp. nov., isolated from olive knots caused by Pseudomonas savastanoi pv. savastanoi. Int J Syst Evol Microbiol 2011;61(Pt 11):2745–2752.

26. Hacker J, Kaper JB. Pathogenicity islands and the evolution of microbes. Annu Rev Microbiol 2000;54:641–679.

27. Messerer M, Fischer W, Schubert S. Investigation of horizontal gene transfer of pathogenicity islands in Escherichia coli using next-generation sequencing. PLoS One 2017;12(7):e0179880.

28. Schneider G, Dobrindt U, Middendorf B, Hochhut B, Szijarto V et al. Mobilisation and remobilisation of a large archetypal pathogenicity island of uropathogenic Escherichia coli in vitro support the role of conjugation for horizontal transfer of genomic islands. BMC Microbiol 2011;11:210.

29. Chen YT, Lai YC, Tan MC, Hsieh LY, Wang JT et al. Prevalence and characteristics of pks genotoxin gene cluster-positive clinical *Klebsiella* pneumoniae isolates in Taiwan. Sci Rep 2017;7:43120.

30. Dubois D, Delmas J, Cady A, Robin F, Sivignon A et al. Cyclomodulins in urosepsis strains of Escherichia coli. J Clin Microbiol 2010;48(6):2122–2129.

31. Johnson JR, Johnston B, Kuskowski MA, Nougayrede JP, Oswald E. Molecular epidemiology and phylogenetic distribution of the Escherichia coli pks genomic island. J Clin Microbiol 2008;46(12):3906–3911.

32. Lai YC, Lin AC, Chiang MK, Dai YH, Hsu CC et al. Genotoxic *Klebsiella* pneumoniae in Taiwan. PLoS One 2014;9(5):e96292.

33. Micenkova L, Benova A, Frankovicova L, Bosak J, Vrba M et al. Human Escherichia coli isolates from hemocultures: Septicemia linked to urogenital tract infections is caused by isolates harboring more virulence genes than bacteraemia linked to other conditions. Int J Med Microbiol 2017;307(3):182–189.

34. Nowrouzian FL, Oswald E. Escherichia coli strains with the capacity for long-term persistence in the bowel microbiota carry the potentially genotoxic pks island. Microb Pathog, 2012;53(3-4):180–182.

35. Krieger JN, Dobrindt U, Riley DE, Oswald E. Acute Escherichia coli prostatitis in previously health young men: bacterial virulence factors, antimicrobial resistance, and clinical outcomes. Urology, 2011;77(6):1420–1425.

36. McCarthy AJ, Martin P, Cloup E, Stabler RA, Oswald E et al. The genotoxin colibactin is a determinant of virulence in Escherichia coli K1 experimental neonatal systemic infection. Infect Immun, 2015;83(9):3704–3711.

37. Yoshikawa Y, Tsunematsu Y, Matsuzaki N, Hirayama Y, Higashiguchi F et al. Characterization of colibactin-producing Escherichia coli isolated from Japanese patients with colorectal cancer. Jpn J Infect Dis 2020.

38. Lam MMC, Wick RR, Wyres KL, Gorrie CL, Judd LM et al. Genetic diversity, mobilisation and spread of the yersiniabactin-encoding mobile element ICEKp in *Klebsiella* pneumoniae populations. Microb Genom 2018;4(9).

39. Shen P, Berglund B, Chen Y, Zhou Y, Xiao T et al. Hypervirulence Markers Among Non-ST11 Strains of Carbapenem- and Multidrug-Resistant *Klebsiella* pneumoniae Isolated From Patients With Bloodstream Infections. Front Microbiol 2020;11:1199.

40. Bankevich A, Nurk S, Antipov D, Gurevich AA, Dvorkin M et al. SPAdes: a new genome assembly algorithm and its applications to single-cell sequencing. J Comput Biol 2012;19(5):455–477.

41. Seemann T. Prokka: rapid prokaryotic genome annotation. Bioinformatics, 2014;30(14):2068–2069.

42. Camacho C, Coulouris G, Avagyan V, Ma N, Papadopoulos J et al. BLAST+: architecture and applications. BMC Bioinformatics 2009;10:421.

43. Blin K, Pascal Andreu V, de Los Santos ELC, Del Carratore F, Lee SY et al. The antiSMASH database version 2: a comprehensive resource on secondary metabolite biosynthetic gene clusters. Nucleic Acids Res 2019;47(D1):D625–D630.

44. Assefa S, Keane TM, Otto TD, Newbold C, Berriman M. ABACAS: algorithm-based automatic contiguation of assembled sequences. Bioinformatics 2009;25(15):1968–1969.

45. Lassmann T, Sonnhammer EL. Kalign - an accurate and fast multiple sequence alignment algorithm. BMC Bioinformatics 2005;6:298.

46. Croucher NJ, Page AJ, Connor TR, Delaney AJ, Keane JA et al. Rapid phylogenetic analysis of large samples of recombinant bacterial whole genome sequences using Gubbins. Nucleic Acids Res 2015;43(3):e15.

47. Stamatakis A. RAxML version 8: a tool for phylogenetic analysis and post-analysis of large phylogenies. Bioinformatics 2014;30(9):1312–1313.

48. Benson G. Tandem repeats finder: a program to analyze DNA sequences. Nucleic Acids Res 1999;27(2):573–580.

49. Bian X, Fu J, Plaza A, Herrmann J, Pistorius D et al. In vivo evidence for a prodrug activation mechanism during colibactin maturation. Chembiochem 2013;14(10):1194–1197.

50. Morgan RN, Saleh SE, Farrag HA, Aboulwafa MM. Prevalence and pathologic effects of colibactin and cytotoxic necrotizing factor-1 (Cnf 1) in Escherichia coli: experimental and bioinformatics analyses. Gut Pathog 2019;11:22.

51. Naughton LM, Romano S, O’Gara F, Dobson ADW. Identification of Secondary Metabolite Gene Clusters in the Pseudovibrio Genus Reveals Encouraging Biosynthetic Potential toward the Production of Novel Bioactive Compounds. Front Microbiol 2017;8:1494.

52. Zha L, Jiang Y, Henke MT, Wilson MR, Wang JX et al. Colibactin assembly line enzymes use S-adenosylmethionine to build a cyclopropane ring. Nat Chem Biol 2017;13(10):1063–1065.

53. Zhang H, Wang Y, Pfeifer BA. Bacterial hosts for natural product production. Mol Pharm 2008;5(2):212–225.

