## Supplemental Figure 1 for "Diversity and prevalence of colibactin- and yersiniabactin encoding mobile genetic elements in enterobacterial populations: insights into evolution and co-existence of two bacterial secondary metabolite determinants"

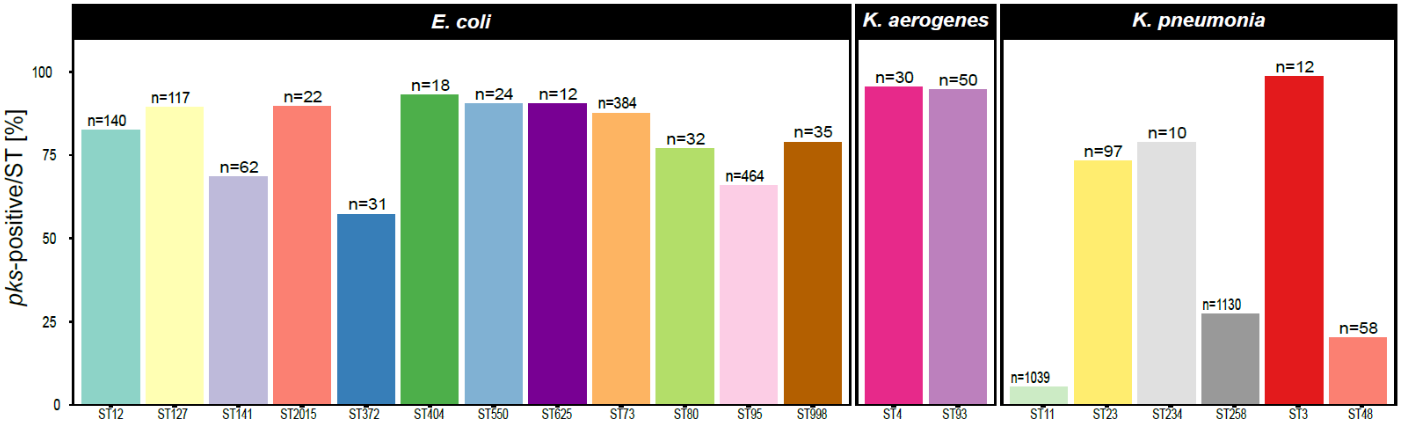


**Figure S1.** Sequence type distribution of *clb*-positive *E. coli* and *Klebsiella* strains.

We compared the amount of strains identified to harbor the *clb* gene cluster with the total number of strains belonging to their respective sequence types in our strain set. The absolute number of strains belonging to sequence types is shown on top of the bars.
