## Supplemental Figure 2 for "Diversity and prevalence of colibactin- and yersiniabactin encoding mobile genetic elements in enterobacterial populations: insights into evolution and co-existence of two bacterial secondary metabolite determinants"

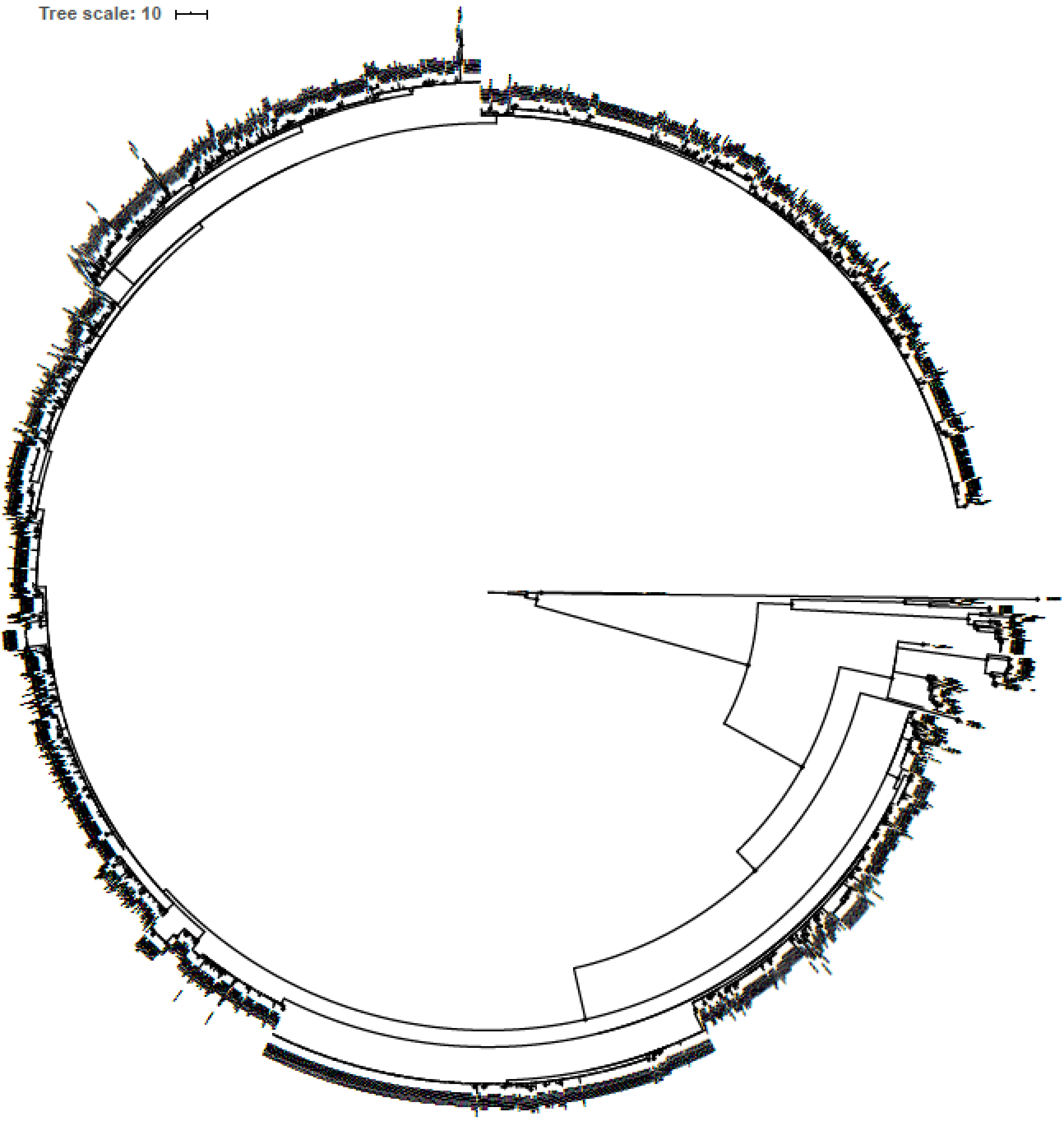


**Figure S2.** Maximum-likelihood based phylogeny of the colibactin gene cluster.

This phylogenetic tree represents all detected colibactin gene clusters in 2169 enterobacterial genomes (included also in Fig. 2A and Fig. 3), including strain labels and branch support values.
