## Supplemental Figure 3 for "Diversity and prevalence of colibactin- and yersiniabactin encoding mobile genetic elements in enterobacterial populations: insights into evolution and co-existence of two bacterial secondary metabolite determinants"

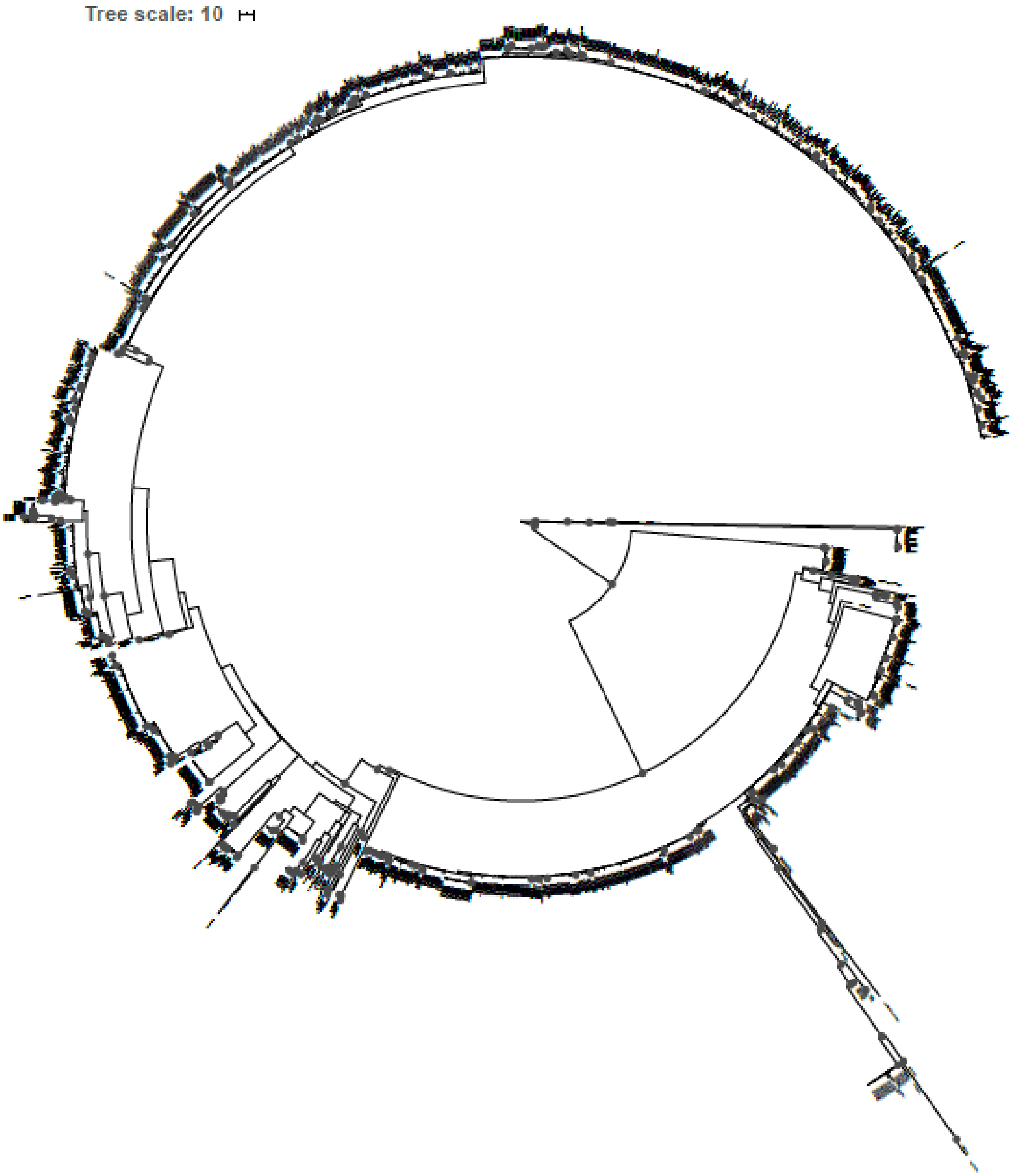


**Figure S3.** Maximum-likelihood based phylogeny of the yersiniabactin gene cluster.
This phylogenetic tree represents all yersiniabactin gene clusters as detected in colibactin positive enterobacterial genomes (included also in Fig. 2B), including strain labels and branch support values.
