## Supplemental Figure 4 for "Diversity and prevalence of colibactin- and yersiniabactin encoding mobile genetic elements in enterobacterial populations: insights into evolution and co-existence of two bacterial secondary metabolite determinants"

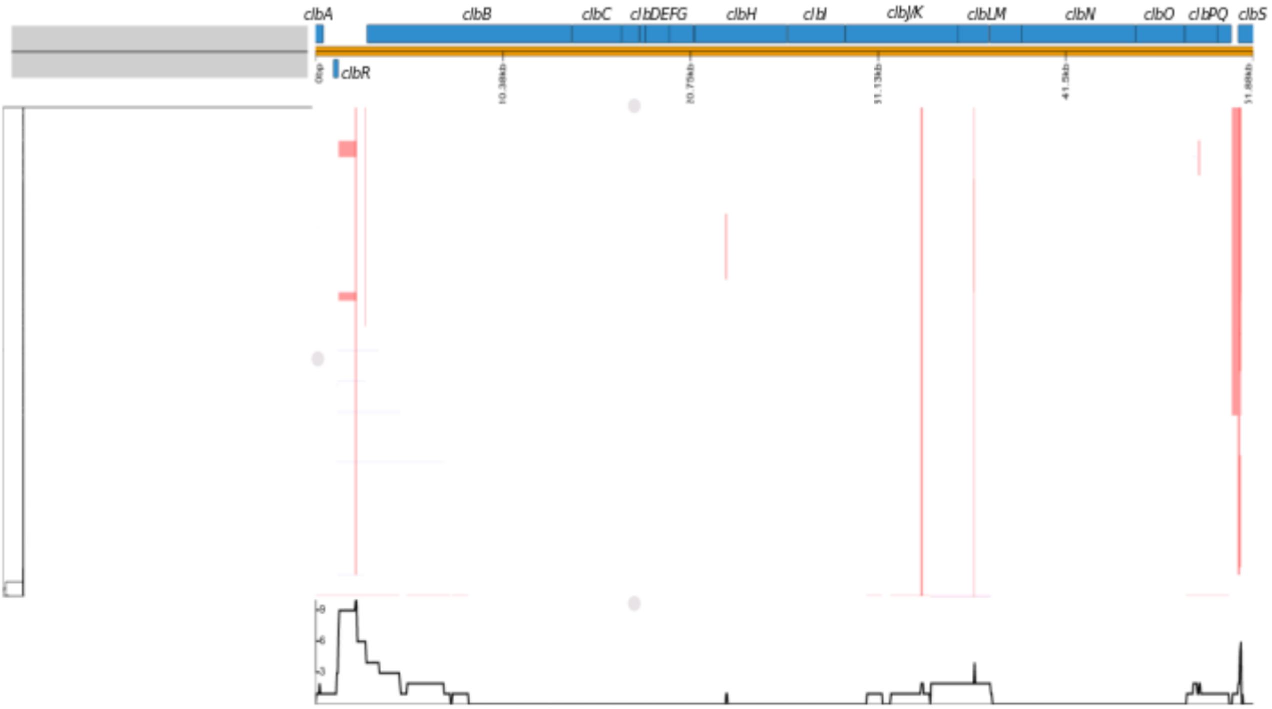


**Figure S4.** Predicted recombination events within the colibactin gene cluster.

Recombination events predicted by Gubbins within the colibactin gene cluster, alongside the phylogeny as shown on the left. The colibactin genes found in the reference genome used (*E. coli* strain M1/5) are shown with blue blocks. The predicted sites of recombination are shown in red blocks.
