## Supplemental Figure 5 for "Diversity and prevalence of colibactin- and yersiniabactin encoding mobile genetic elements in enterobacterial populations: insights into evolution and co-existence of two bacterial secondary metabolite determinants"

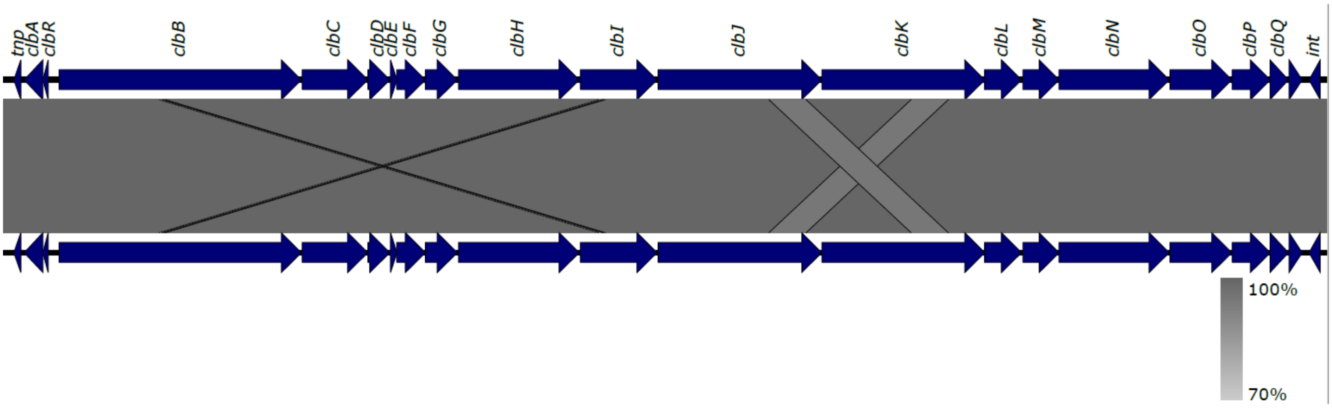


**Figure S5.** Linear comparison of the nucleotide sequence of the colibactin gene cluster.

Comparison of the colibactin genes found in *E. coli* strain M1/5 against themselves using Easyfig (Sullivan et al. 2011). The alignment of colibactin genes, with a minimum nucleotide identity of 70%, shows the homology between regions of *clbJ* and *clbK*, which was discussed to be a potential cause for misassembly in short-read sequencing data.
