## Supplemental Figure 6 for "Diversity and prevalence of colibactin- and yersiniabactin encoding mobile genetic elements in enterobacterial populations: insights into evolution and co-existence of two bacterial secondary metabolite determinants"

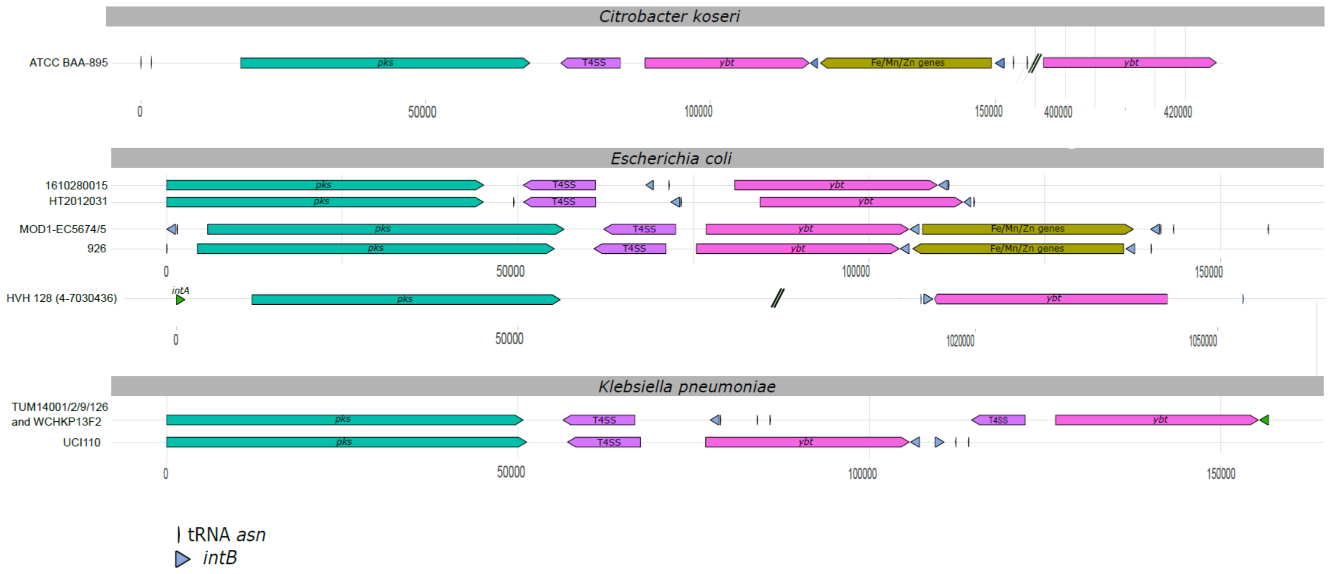


**Figure S6.** Structural variation of the *clb* and *ybt* gene cluster-harboring ICEs in *E. coli*, *K. pneumoniae*, and *C. koseri*.

The different structures found within ICEs that do not conform to the major structures shown in Fig. 4 are depicted. The *clb* gene cluster (teal green), T4SS module (purple), *ybt* determinant (pink), integrase genes (green/blue), and the Fe/Mn/Zn module (yellow) found within the ICE are shown.
