## Supplemental Figure 7 for "Diversity and prevalence of colibactin- and yersiniabactin encoding mobile genetic elements in enterobacterial populations: insights into evolution and co-existence of two bacterial secondary metabolite determinants"

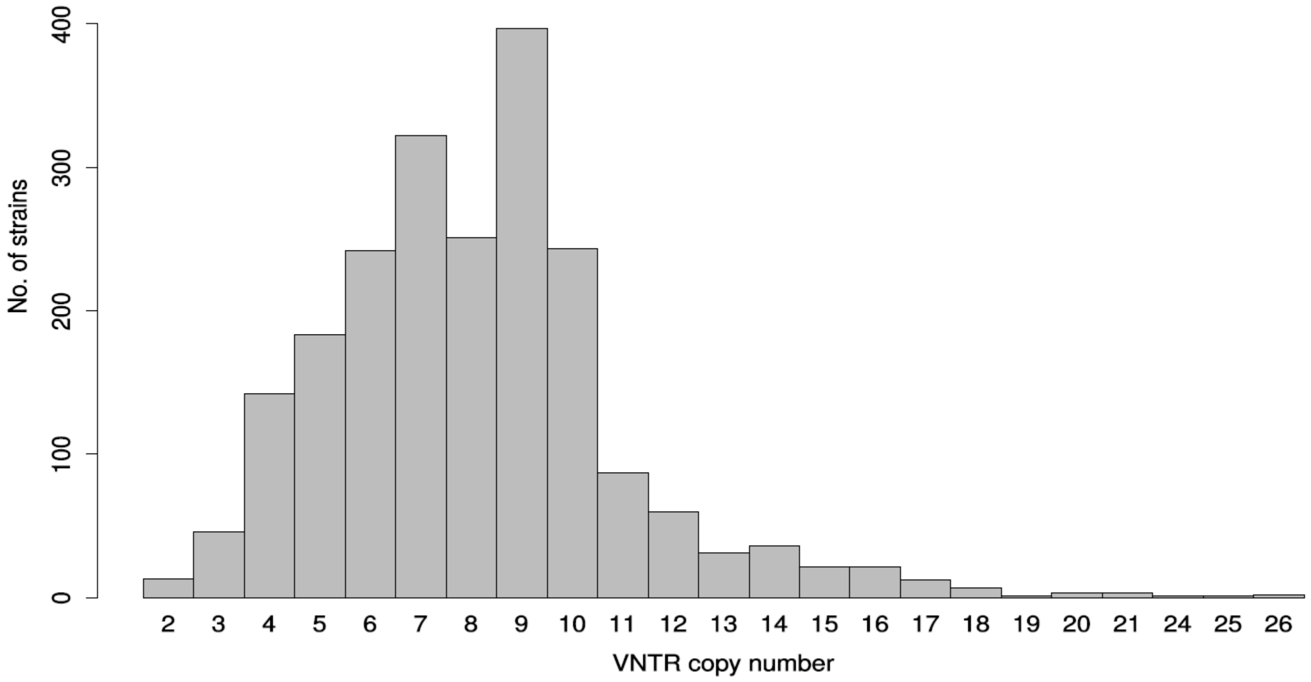


**Figure S7.** Size distribution of the variable number of tandem repeats (VNTR) region in *clb* positive bacterial strains.

The number of repeats detected in the VNTR region located between *clbR* and *clbB* is shown in relation to the frequency of their occurrence in bacterial genomes.
