## Supplemental Figure 8 for "Diversity and prevalence of colibactin- and yersiniabactin encoding mobile genetic elements in enterobacterial populations: insights into evolution and co-existence of two bacterial secondary metabolite determinants"

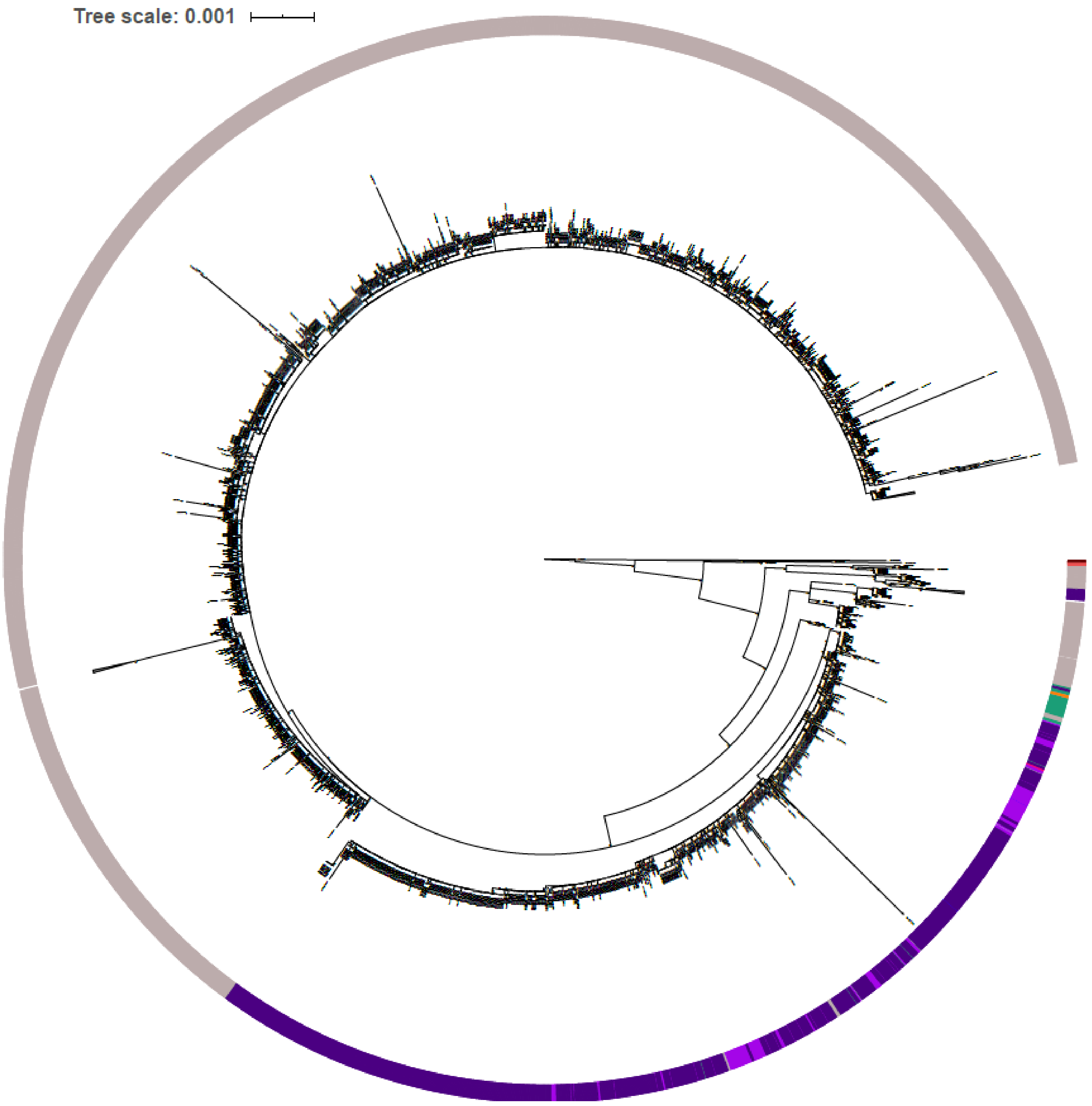


**Figure S8.** Phylogeny inferred from amino acid sequences of 17 *clb* genes. The amino acid sequences from colibactin genes were concatenated and used to generate a phylogeny in a similar manner to that of the nucleotide-based phylogeny. The outer circle describes the species of each node/strain harboring the colibactin genes.
